# Molecular mechanisms reconstruction from single-cell multi-omics data with HuMMuS

**DOI:** 10.1101/2023.06.09.543828

**Authors:** Remi Trimbour, Ina Maria Deutschmann, Laura Cantini

## Abstract

The molecular identity of a cell results from a complex interplay between heterogeneous molecular layers. Recent advances in single-cell sequencing technologies have opened the possibility to measure such molecular layers of regulation.

Here, we present HuMMuS, a new method for inferring regulatory mechanisms from single-cell multi-omics data. Differently from the state-of-the-art, HuMMuS captures cooperation between biological macromolecules and can easily include additional layers of molecular regulation.

We benchmarked HuMMuS with respect to the state-of-the-art on both paired and unpaired multi-omics datasets. Our results proved the improvements provided by HuMMus in terms of TF targets, TF binding motifs and regulatory regions prediction. Finally, once applied to snmC-seq, scATAC-seq and scRNA-seq data from mouse brain cortex, HuMMuS enabled to accurately cluster scRNA profiles and to identify potential driver TFs.

## Introduction

Cells within a multicellular organism are remarkably heterogeneous, spanning many different molecular identities^1, 2^. The molecular identity of a cell is the result of a complex interplay among different layers of molecular regulation, all of which can vary because of intrinsic and extrinsic factors. Recent advances in single-cell sequencing technologies have opened the possibility to measure such molecular layers of regulation, a.k.a. omics, at the resolution of the single cell. Examples of omics data currently accessible at single-cell resolution are chromatin accessibility (scATAC), methylation (snmC), expression (scRNA)^3, 4^. In addition, sequencing technologies providing the joint profiling of multiple single-cell omics from the same cell have been developed^5, 6^. Examples of them are 10xGenomics Multiome platform, jointly profiling transcriptome and chromatin accessibility from the same cell, and CITE-seq, simultaneously quantifying cell surface proteins and transcriptome within a single cell^7^. All these data provide the unprecedented opportunity to reveal how different molecular layers interact through complex regulatory mechanisms to define cell identity.

Several methods, co-analysing single-cell omics data to elucidate the regulatory mechanisms that encode cellular identities, have been recently developed^8–14^. The output of these methods are Gene Regulatory Networks (GRNs), corresponding to graphs linking Transcription Factors (TFs) with their inferred target genes and/or peaks^15–17^. The GRNs are obtained by all methods performing TF-peak-gene associations based on binding motif databases (e.g. JASPAR^18^), then filtered through scRNA and scATAC data analysis. All these methods ignore intra-omics cooperation between biological macromolecules, which is crucial in biology. Indeed, TFs can cooperate in the regulation of gene expression by forming dimers and multiple DNA regions can co-regulate the expression of the same gene. In addition, state-of-the-art methods only consider TF-gene interactions present in binding motifs databases and miss all those interactions that are not reported there. Furthermore, all these methods infer GRNs by integrating scRNA and scATAC data, thus ignoring all other complementary layers of molecular regulation (e.g. methylation, proteome). Finally, many methods require either paired data, or perform cell pairing before GRN inference^11–14^. This is a major limitation, as paired single-cell multi-omics data are still rare and performing cell pairing in dataset profiled from different cells forces a decrease in the size of one of the two datasets thus reducing the richness of its information content.

Here we introduce HeterogeneoUs Multilayers for MUlti-omics Single-cell data (HuMMuS), a flexible tool based on Heterogeneous Multilayer Networks (HMLNs) to reconstruct regulatory mechanisms from multiple single-cell omics data. HuMMuS considers not only inter-omics interactions (e.g. peak-gene, TF-peak), as done by the state-of-the-art, but also intra-omics ones (e.g. peak-peak, gene-gene, TF-TF) thus allowing to capture cooperation between biological macromolecules. This inclusion of intra-omics interactions allows HuMMuS to explore new TF-gene interactions not present in binding motif databases. In addition, HuMMuS is a flexible framework, that can be used both for paired and unpaired single-cell multi-omics data or easily extended to deal with additional omics data, thus not limiting the regulatory mechanisms analysis to only scRNA and scATAC, as it is currently done in the state-of-the-art.

We extensively benchmarked HuMMus with respect to the state-of-the-art on four independent datasets of scRNA and scATAC. This benchmarking included the prediction of TF targets, TF binding regions, regulatory regions and the association of its communities with known biological processes. Finally, by applying HuMMuS to unpaired scRNA, scATAC and scnmC data from mouse cortex, we showed that its GRN allows to accurately cluster scRNA profiles and to identify regulators relevant to mouse brain cortex.

HuMMus is available at https://github.com/cantinilab/HuMMuS as R package, together with a tutorial for its usage.

## Results

### HuMMuS a new tool for molecular mechanisms reconstruction from single-cell multi-omics data

We developed HeterogeneoUs Multilayers for MUlti-omics Single-cell data (HuMMuS), a new tool for regulatory mechanisms inference from single-cell multi-omics data (Figure 1, https://github.com/cantinilab/HuMMuS).

**Figure 1.**
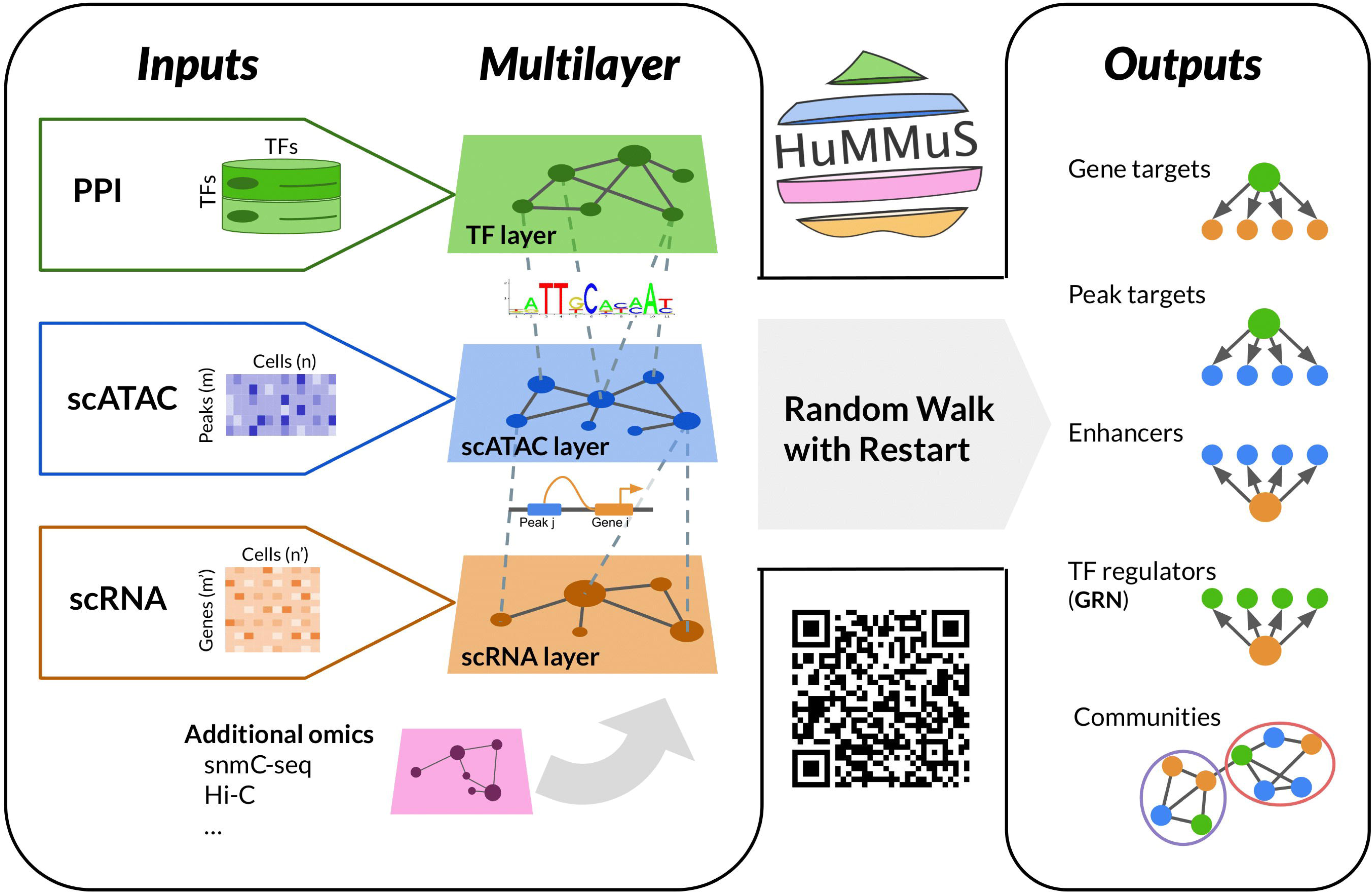
Schematic view of HuMMuS workflow.

HuMMuS is based on Heterogeneous Multilayer Networks (HMLNs). A HMLN is a network *M* = (*V_m_*, *E_m_*, **L**), *m* = 1, …, *M*, composed of *M*, layers each of them containing different nodes and different intra-layer links *E_m_* ⊆ *V_m_* × *V_m_*. Nodes of different layers are connected by inter-layers links encoded in **L**^19, 20^. As summarized in Figure 1, we reconstruct HMLNs composed of three layers: The TF layer, containing unlinked TFs, the scATAC layer containing peak co-accessibility information inferred from scATAC data and the scRNA layer encoding transcriptional regulation inferred from scRNA data. For all details on the layers’ construction see Methods. Of note, we here focused on this combination of omics data to not advantage HuMMuS by the additional information provided by other single-cell omics data. However, as the HMLN structure is flexible, HuMMuS can easily integrate other single-cell omics data, such as methylation (snmC) or Hi-C data, and additional information on known interactions, such as Protein-Protein interactions in the TF layer to capture TFs cooperativity. Once the HMLN is constructed, HuMMuS uses Random Walks with Restart (RWR)^20^ to mine the HMLN and extract different outputs: (i) the prediction of the targets of a Transcription Factor (TF), based on RWRs starting from each TF in the TF layer and exploring the full network until the scRNA layer; (ii) the prediction of the peaks bound by a given TF, based on RWRs starting from each TF in the TF layer and exploring the scATAC layer; (iii) the prediction of the regulatory regions (proximal and distal enhancers) associated to a given gene, based on RWRs starting in each gene of the scRNA layer and exploring the scATAC layer; (iv) the reconstruction of Gene Regulatory Networks (GRNs), based on RWRs starting in each gene of the scRNA layer and exploring the full network until the TF layer; (v) the extraction of communities in the GRN, reflecting tightly connected macromolecules in the HMLN frequently involved in the regulation of the same biological process or pathway^21^. Of note, both the prediction of TF targets (output i) and the reconstruction of the GRNs (output iv), in principle lead to a TF-gene network. The choice of reconstructing GRNs by exploring the HMLN from genes to TFs is justified by the need of having a competition among different TFs in the regulation of a gene, as done in most of the GRN inference approaches^8–17^. On the contrary, when predicting the targets of a TF, we want to treat each TF independently from the others and make genes compete among themselves. For this reason, we obtain the output (i) by exploring the HMLN from TFs to genes. See methods for all details on the parameter choice for the RWR and the possible outputs.

Thanks to the use of a HMLN structure, HuMMuS has multiple advantages with respect to the state-of-the-art. First, it captures not only inter-omics interaction (e.g. peak-gene, TF-peak), as done by the state-of-the-art, but also intra-omics ones (e.g. peak-peak, gene-gene, TF-TF). This allows HuMMuS to capture cooperation between biological macromolecules and use it to predict, for example, TF-gene interactions not present in binding motifs databases. In addition, HuMMuS is a flexible framework, that can be used both for paired and unpaired single-cell multi-omics data or easily extended to deal with additional omics data, thus not limiting the regulatory mechanisms analysis to only scRNA and scATAC, as it is currently done in the state-of-the-art.

In the following we extensively benchmark HuMMuS against CellOracle and Pando^10, 11^, being the most famous published works in the field. Interestingly, CellOracle is the only existing method considering some cooperation at the peaks level. In addition, we included GENIE3 in the benchmark as a baseline for performances when considering scRNA alone. All the benchmarking is performed on four test cases (see Methods and Supp Table 1): two datasets (called in the following Chen and Liu) of human Embryonic Stem Cells (hESCs), jointly profiled for scRNA and scATAC (i.e. paired data), and two unpaired scRNA and scATAC datasets of mouse Embryonic Stem Cells (mESCs) (called in the following Düren and Semrau). For details on HuMMuS layers structure in these four datasets see Supp Table 2. Of note, in Düren and Semrau, being the data unpaired, the scRNA and scATAC information has been profiled from different cells all extracted from mESCs. These last two test cases thus allow to test the impact of cell pairing on the performances of the different methods. The choice of these four test cases is justified by the availability of ChiP-seq and Transcription Factor perturbation experiments in hESCs and mESCs from^17^. These additional data, already used in benchmarking works^17^, allow indeed to build good ground truths for the different tests presented in the following sections.

### HuMMus outperforms the state-of-the-art in Transcription Factor (TF) target prediction

We first focused on benchmarking HuMMuS with respect to the state-of-the-art based on the quality of its Transcription Factor (TF) targets predictions. This analysis has been performed on the four test cases presented above, corresponding to scRNA and scATAC profiling of hESCs and mESCs. As ground truth of the TF-targets interactions we used the intersection between ChIP-seq and TF perturbations experiments, as done in^17^. This choice represents indeed the best estimation of TF targets we can get for real data, as it assures the presence of a binding site for the TF on the promoter of the target gene and, at the same time, a downregulation of the target gene once the TF is knocked down/out.

As described in Figure 2A, in each of the four test cases, HuMMus and the other state-of-art algorithms have been independently applied, a ranking of putative targets for each TF is then identified and compared with the ground truth described above. The ranking of putative gene targets for a TF is obtained for the state-of-the art methods as the list of genes linked to the TF. The genes are ordered according to the weight of their links. For HuMMuS instead, we perform a Random Walk with Restart (RWR) starting from each TF and going across all the HMLN, thus obtaining a ranking of putative target genes based on their closeness to the TF. The overlap for all methods with the ground truth is then analyzed when cutting the ranking at different levels (3, 5, 10, 15, 20, 30, 40, 50, 75, 100).

**Figure 2.**
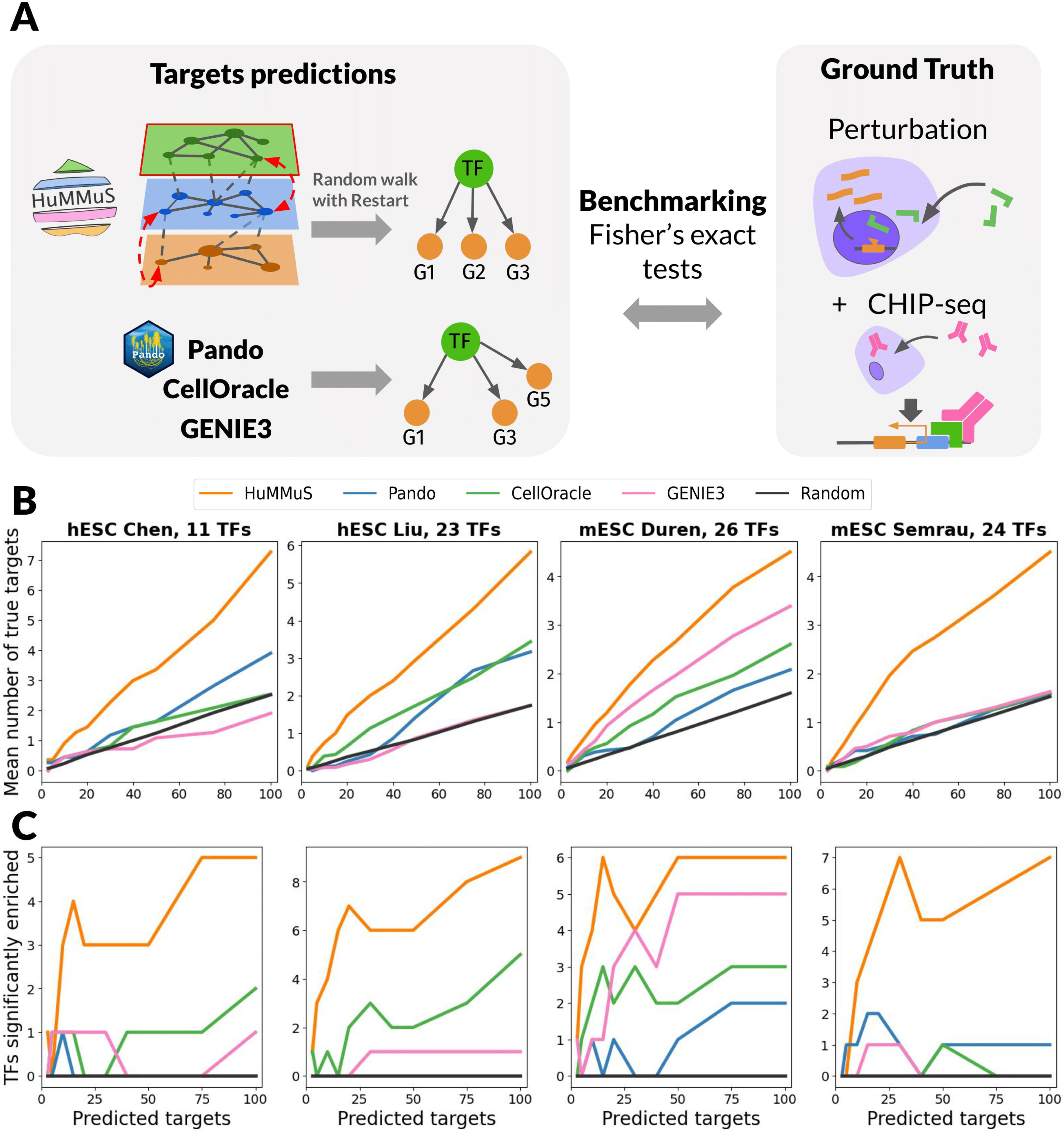
Transcription Factor (TF) targets prediction benchmarking. (A) schematic view of the performed benchmarking. (B) average number of correctly predicted targets per TF. (C) number of TFs having a significant amount of correctly predicted targets (Fisher’s test p-value <0.05). In (B-C) different colors correspond to different methods: orange (HuMMuS), blue (Pando), green (CellOracle), pink (GENIE3) and black (random).

As shown in Figure 2B, in all four studied test cases HuMMus obtains the highest number of correctly predicted average targets per TF. Of note, in Semrau the results of state-of-the-art methods are close to random, here represented with a black curve. Of note, even when pairing the cells in the two unpaired datasets, the performances observed for HuMMuS are not affected (see Supp Figure 1). To then test whether the observed performances were driven by a subgroup of TFs or consistent for a high number of them, we computed the number of TFs having a significant number of targets in their top predicted targets (see Methods for details). As shown in Figure 2C, overall, all methods get few TFs with a significant amount of correctly predicted targets. At the same time, also in this case, HuMMus gets best performances in all four test cases. Taken together these two results suggest a high potential for HuMMus in TF targets prediction.

### HuMMuS outperforms the state-of-the-art in regulatory region identification

We then benchmarked HuMMuS with respect to the state-of-the-art based on known regulatory regions identification. This benchmark was realized in two steps: first, the ability to predict the peaks bound by a TF is tested; then, the quality of the regulatory regions (proximal and distal enhancers) predicted for each gene is evaluated. As GENIE3 does not provide any information on regulatory regions, it was excluded from this part of the benchmarking.

As shown in Figure 3A, to test the quality of the peaks associated with a TF, in HuMMuS we used RWRs from each TF as a proxy of the compatibility between a TF and peaks and filtered the obtained peak ranking at different levels (100%, 80%, 60%, 20%). For CellOracle and Pando instead, we considered the peaks retained by the model as associated with each TF (see Methods for details). In CellOracle different peak co-accessibility correlation thresholds have been considered 0.05, 0.2 and 0.8, with the last being the default threshold. We finally compared the predictions obtained by the various methods with the ground-truth composed of ChiP-seq experiments results on the biological system under analysis (mESCs and hESCs) from^22^. See methods for further details on the analysis.

**Figure 3.**
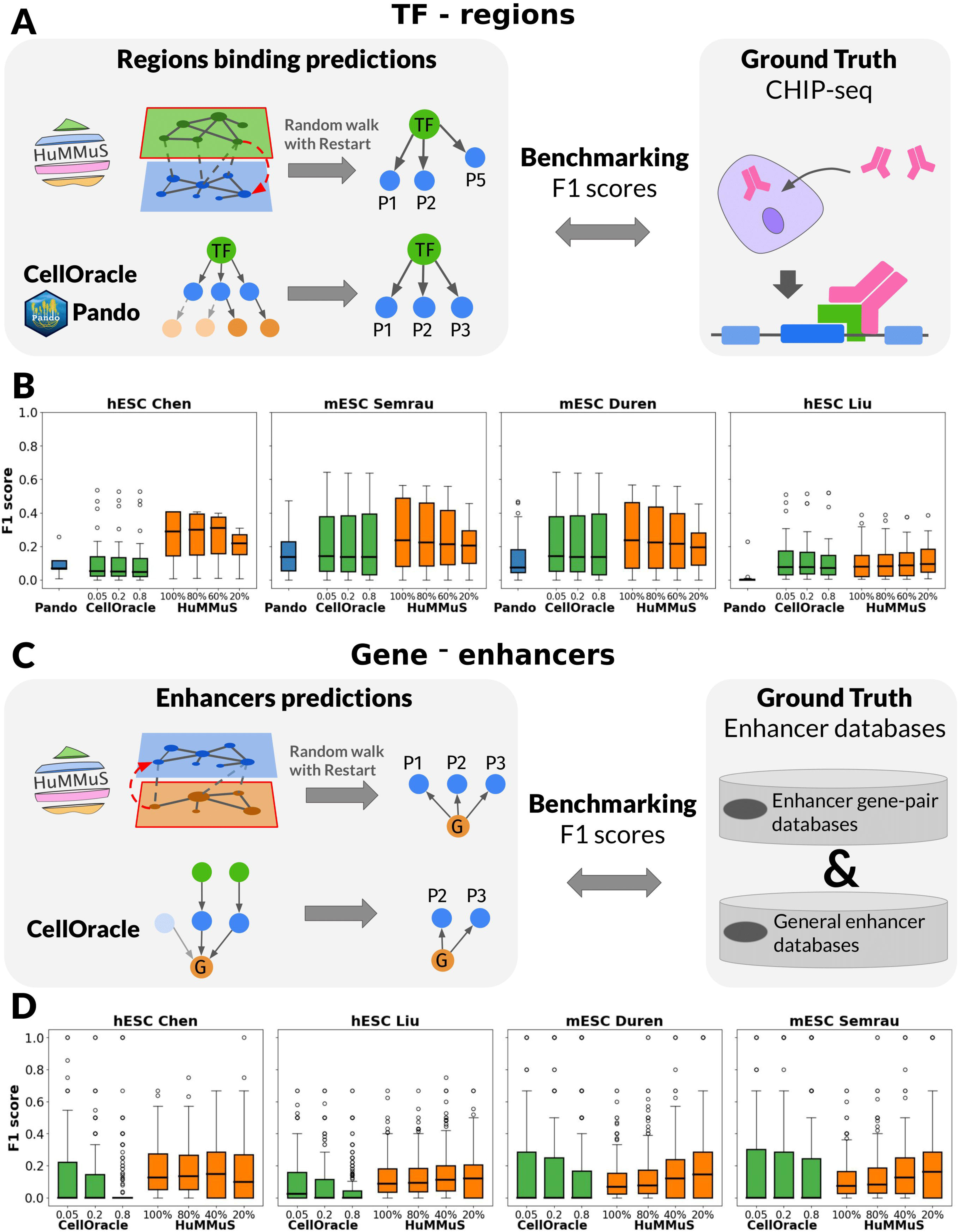
Regulatory regions benchmarking. (A) schematic view of the benchmarking performed for TF-peak associations. (B) F1 score of the intersection between the ground-truth TF-peak associations and those inferred by Pando, CellOracle and HuMMuS; the 100%, 80%, 60%, 20% thresholds of HuMMuS correspond to the number of nodes retained from the RWRs ranking. For CellOracle instead, 0.05, 0.2 and 0.8 correspond to the correlation thresholds of the model, with 0.8 being the default one. (C) schematic view of the benchmarking performed for gene-peak associations. (D) F1 score of the intersection between the ground-truth gene-peak associations and those inferred by CellOracle and HuMMuS. In (B,D) different colors correspond to different methods: orange (HuMMuS), blue (Pando), green (CellOracle). The thresholds are the same as those of panel (B).

Overall, as shown in Supp Figure 2A, HuMMuS identifies more peaks associated with a TF than alternative methods. This result is not surprising as, differently from the state-of-the-art, HuMMuS leverages all the peak layer without constraints neither on genomic windows neither on known TF motifs. More interestingly, as shown in Figure 3B, once checking the quality of the identified TF-peak associations, HuMMuS shows higher F1 scores for all the considered thresholds.

We then focused on the regulatory regions associated with each gene. As shown in Figure 3C, in HuMMuS we predicted the peaks having a regulatory role on a gene based on RWRs starting from the gene and filtered the obtained ranking at (100%, 80%, 60%, 20%). For CellOracle instead, the peaks associated to a gene by its model were considered and filtered with different correlation thresholds: 0.05, 0.2 and 0.8, with the last being the default one. The obtained predictions were finally compared with a ground truth composed of gene-regulatory regions associations available from different databases^23–29^. For all details on the analysis, see Methods. GENIE3 and Pando have been excluded from this analysis as they did not provide an output allowing for this type of evaluation.

As shown in Supp Figure 2B, overall HuMMuS gets more enhancers associated with each gene. Again, this result is not surprising given that the intrinsic structure of HuMMuS allows it to predict new peak-gene associations, without genomic windows constraints. In addition, as shown in Figure 3D HuMMuS has a higher F1 score than the state-of-the-art, indicating that the regulatory regions predicted by HuMMuS tend to more frequently reflect known ones. In addition, HuMMuS shows an overall improvement of the F1 score when keeping only the highest scored predicted enhancers. This suggests that the scores provided by HuMMuS provide a meaningful ranking of the potential enhancers. On the contrary, CellOracle shows a decrease in performance once increasing the peak co-accessibility correlation threshold.

Taken together these two results suggest that HuMMuS can powerfully predict regulatory regions associated with TF gene regulation. Also in this case, the results observed for HuMMuS in the two unpaired data (Duren and Semrau) are not affected by cell pairing (Supp Figure 3).

### HuMMuS outperforms the state-of-the-art in the biological relevance of its gene communities

We finally benchmarked HuMMuS with respect to the state-of-the-art based on the biological relevance of their gene communities. Indeed, gene communities in biological graphs have been previously shown to frequently reflect known pathways and biological processes^21, 30, 31^.

As shown in Figure 4A, the Louvain algorithm^32^ was applied to the HuMMuS GRN and to those of the state-of-the-art and the biological relevance of the obtained communities was evaluated based on the percentage of communities enriched in pathways (KEGG^33, 34^ and REACTOME^35^) and Gene Ontologies^36, 37^. Before running community detection, as most of the GRNs are highly dense (density >0.8 in half of networks see Supp Table 3), a filtering was applied to the links to make all networks equally dense. Regarding the community detection, as the Louvain algorithm depends on the resolution parameter, we here run it with resolution varying in the range 0-2 and choose for each method the resolution giving best performances and a reasonable number of communities (≥ 10). See Methods for details on the analysis, Supp Table 4 for performances across different resolution values.

**Figure 4.**
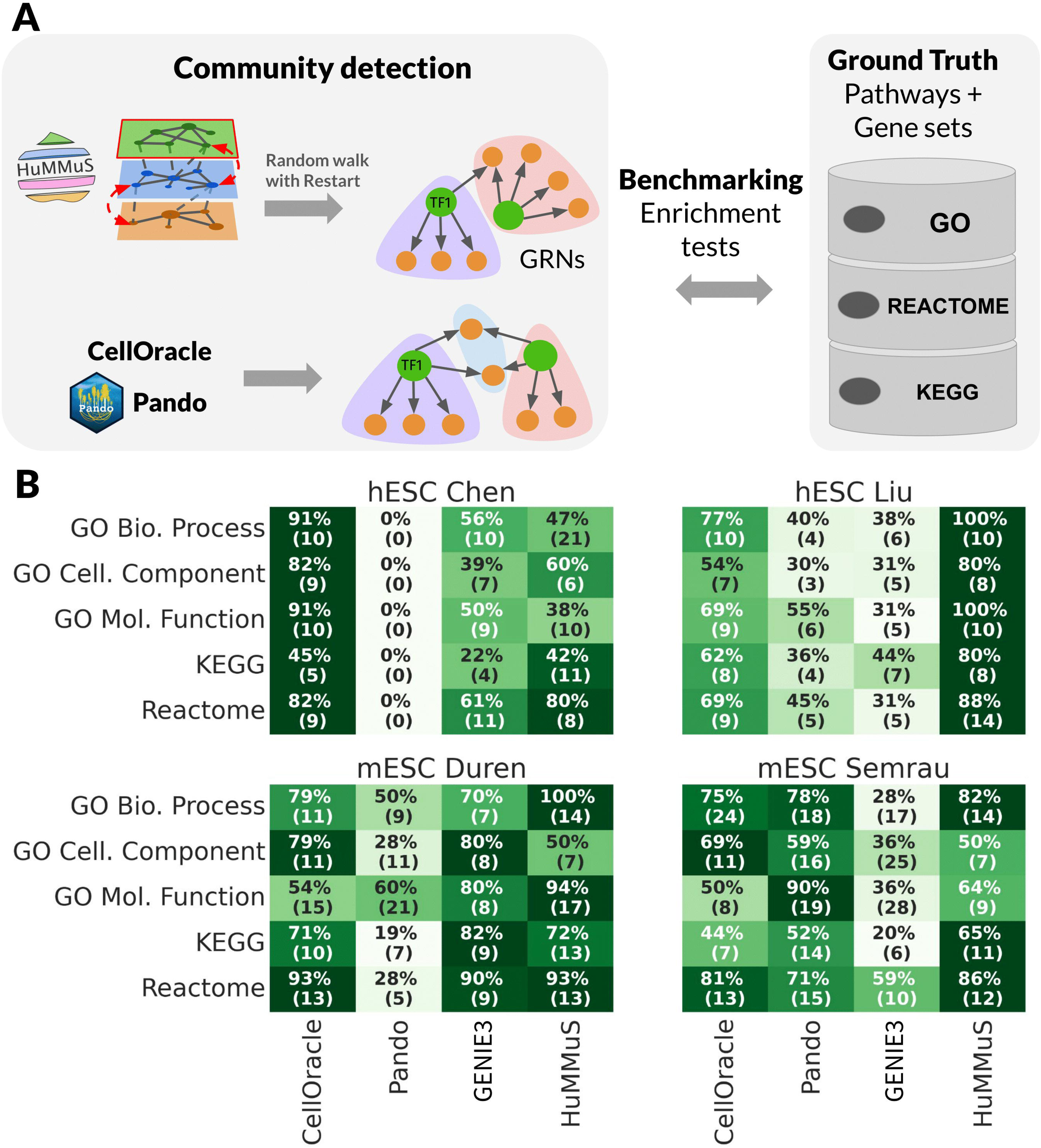
Community detection benchmarking. (A) schematic view of the benchmarking performed for community detection. (B) heatmaps of percentage of enriched community in each benchmarked method across the five biological databases. The values reported in the table correspond to the percentage of enriched communities, while those in parentheses are the actual number of enriched communities.

Figure 4B shows the results of the comparison. Regarding the number of communities corresponding to the best enrichment performances, all methods vary in a range of 10-30 communities, depending on the test case and the database under analysis. Concerning the enrichment in pathways and Gene Ontologies, in three out of four test cases (Liu, Duren and Semrau), HuMMuS gets the highest percentage of enriched communities in most of the databases. In the remaining test case (Chen), CellOracle gets better results. Of note, no evident correlation emerges between the number of identified communities and the performances of the different methods (see Supp Table 4).

### Challenging HuMMus in mouse cortex profiled for scRNA, scATAC and scnmC

We finally challenged HuMMuS in the reconstruction of molecular mechanisms of the mouse brain cortex. Differently from the state-of-the-art, here for the first time we take into account three single-cell omics data: scRNA^38^, scATAC^39^ and scmC^40^. The data of size 55,803 cells in scRNA, 2317 cells in scATAC and 3386 cells in scnmC are unpaired, obtained by profiling mouse cortical neurons.

Following the HuMMuS pipeline, we reconstructed a HMLN composed of four layers: TF layer, scATAC layer, scnmC layer and scRNA layer (see Figure 5A). Then RWRs from the scRNA layer have been used to extract a GRN composed of 637 regulons, each corresponding to a TF and its associated genes ranked by the strength of association^41^.

**Figure 5.**
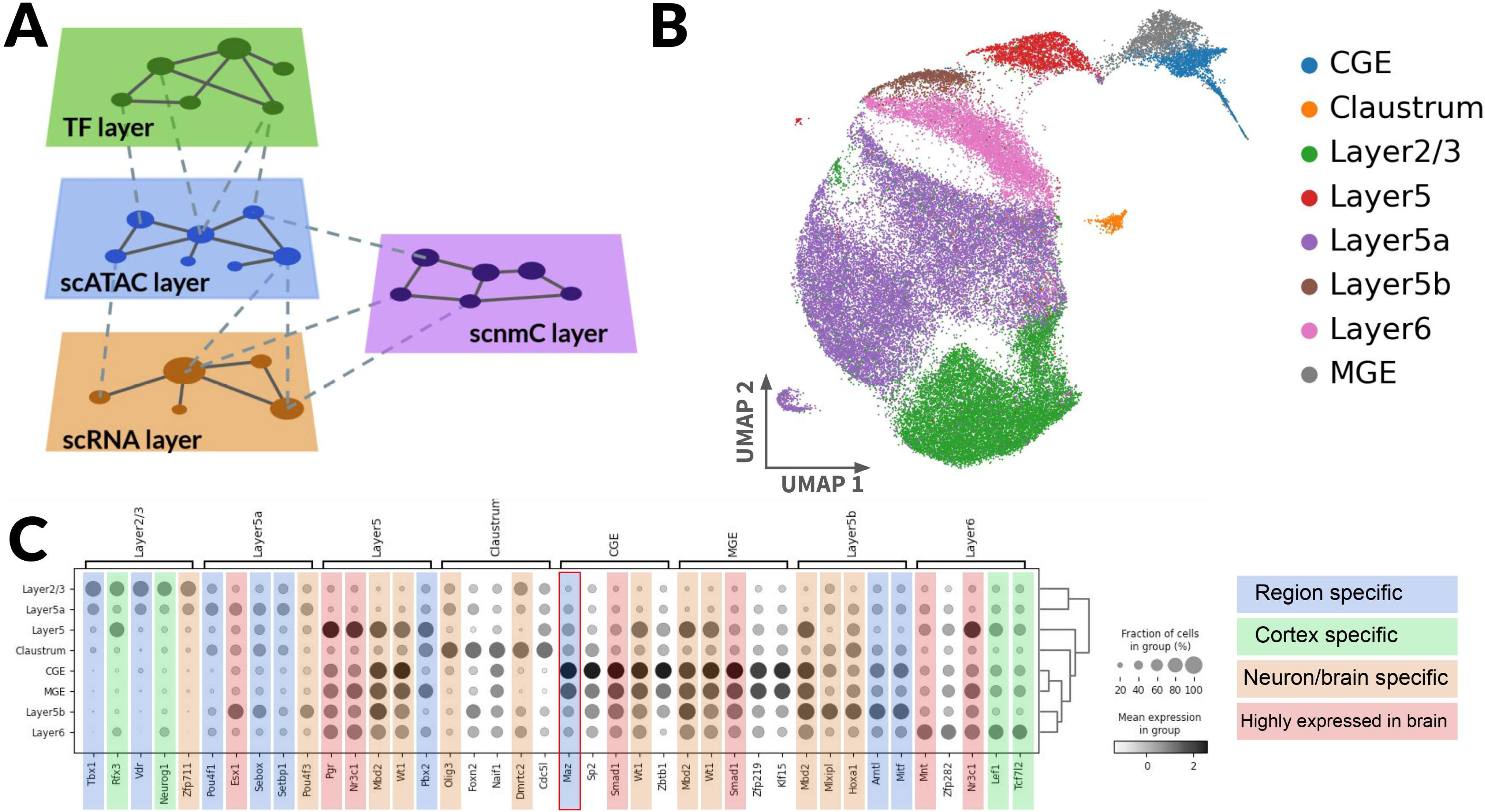
challenging HuMMuS on scRNa, scATAC and scnmC from mouse cortex. (A) HMLN used in HuMMuS to reconstruct regulatory mechanisms from scRNA, scATAC and scnmC. (B) UMAP plot obtained from HuMMuS regulon activity. Cells are colored according to the labels present in their original publication and in previous analyses^38, 43^. (C) Heatmap of activitys

As a first observation, the activity of the obtained regulons, computed according to^41, 42^, is able to correctly cluster the cells according to their area of origin in the mouse cortex (see Figure 5B). This suggests that the regulons identified by HuMMuS can nicely recapitulate the known heterogeneity present between the analyzed cells and already reported in^38, 43^.

We then focused on the regulons strongly associated with each of these cell populations, considering only the top five differentially active regulons per cell population (Figure 5C, Methods for details). Of the obtained 34 regulons, 76% of their TFs have an already reported association with either neurons, cortex, or brain (see Supp Table 5). In particular, five of them (Esx1, Pgr, Nr3C1, Smad1/5, Mnt) are reported in the Bgee database as expressed in the brain^44^. Nine of them (Zfp711, Pou4f3^45^, Mbd2^46^, Wt1^47^, Olig3^48^, Dmrtc2^49^, Mlxipl^50^, Hoxa1^51^) are documented in publications associating them with either brain or neurons and thirteen of them (Tbx1/Tbx10^52^, Rfx3^8, 53^, Neurog1^54^, Vdr^55^, Pou4f1/Pou4f2^56^, Sebox^57^, Setbp1^58^, Pbx2/Pbx4^59^, Maz^60, 61^, Arntl^62^, Mitf^63^, Lef1^64^, Tcf7l2^64^) are reported in publications specifically referring to the mouse cortex. Of note, four of these TFs were also already documented to be associated to the specific region of the cortex where HuMMuS found them to be differentially active. This is the case for Rfx3 and Neurog1, that we find associated with Layer 2/3 and that had been previously associated with this exact brain region^8, 53, 54, 65^. In addition, Lef1 and Tcf17l2 have been documented to be associated with deep layers of the cortex and HuMMuS identifies them in layer 6^64^.

Finally, HuMMuS suggests the possible regulatory role of MAZ into CGE-derived cortical inhibitory interneurons. Through bibliographic research MAZ is documented to have a role in neuronal stem cells differentiation and as potential regulator in Purkinje cells, a gaba-ergic inhibitory neuron population^60, 61^. HuMMuS associates it to the Caudal Ganglionic Eminence (CGE) region, producing a high proportion of cortical inhibitory neurons (30%)^66^. In addition, in the top 10% of the 9341 inferred targets of MAZ, we can find Cntnap3, Dlx5, Sp9, Dlx6, Nr2c2ap, Dlx2, Arx, Grik3, all genes documented to be differentially expressed in inhibitory interneurons in The Mouse Organogenesis Atlas (MOCA)^67^.

## Discussion

Cell identities result from the joint activity of different molecular layers of regulation. These molecular layers can be nowadays measured thanks to single-cell sequencing technologies, such as scRNA, scATAC, scnmC.

Different methods have been recently designed to reconstruct molecular mechanisms from different single-cell omics data. Here we proposed HeterogeneoUs Multilayers for MUlti-omics Single-cell data (HuMMuS), a flexible tool based on Heterogeneous Multilayer Networks (HMLNs) to reconstruct regulatory mechanisms from multiple single-cell omics data. HuMMuS is found to have better performance than the state-of-the-art in the prediction of TF targets, TF binding regions, regulatory regions and in the identification of biologically relevant gene communities. Once applied to the integration of scRNA, scATAC and scnmC data profiled from mouse cortex, HuMMuS identified relevant regulatory mechanisms.

Overall, the main advantages of HuMMuS are the ability to capture intra-omis cooperation between biological macromolecules, its flexibility allowing it to easily integrate additional omics or prior information (e.g. pathway databases) and to work with both paired and unpaired data.

For simplicity, we here only explored inter-layer links based on databases. However, such links could be improved in concrete biological applications considering inter-layer links derived from experimental evidence (e.g. resulting from ChiP-seq experiments instead of generalistic motif databases). In addition, cooperation between TFs is not here considered to not favor HuMMuS over other methods in the benchmarking. However, protein-protein interaction data and ChiP-seq data could be included in the TF layer of HuMMuS as a proxy of TF-TF cooperativity. Finally, we here focused on community detection in GRNs to have a comparable output between HuMMuS and the current state-of-the-art. However, HuMMuS could further include in the future methods for community detection in HMLNs, thus allowing to detect cross-omics communities, providing a better picture of the complex interactions driving some biological processes.

## Methods

### HeterogeneoUs Multilayers for MUlti-omics Single-cell data (HuMMuS)

We developed HeterogeneoUs Multilayers for MUlti-omics Single-cell data (HuMMuS), a new tool for regulatory mechanisms inference from single-cell multi-omics data (https://github.com/cantinilab/HuMMuS).

HuMMuS is based on Heterogeneous Multilayer Networks (HMLNs). A HMLN is a network *M* = (*V_m_*, *E_m_*, **L**), *m* = 1, …, *M*, composed of *M* layers each of them containing different nodes *V_m_* and different intra-layer links *E_m_* ⊆ *V_m_* × *V_m_*. Nodes of different layers are connected by inter-layers links encoded in **L**^19, 20^. As summarized in Figure 1, we reconstruct HMLNs composed of three layers: The TF layer, containing unlinked TFs, the scATAC layer containing peak co-accessibility information inferred from scATAC data and the scRNA layer encoding transcriptional regulation inferred from scRNA data. Details on the layers construction are provided below.

#### Heterogeneous Multilayer Network (HMLN) construction

The standard structure we propose for molecular mechanisms reconstruction with HuMMuS is based on scRNA-seq and scATAC-seq data that doesn’t need to be paired.

##### TF layer

TFs expressed in the scRNA data and having a known motif according to JASPAR or cisBP databases^18, 68^ were included in the TF layer. In the presented results, we did not include TF-TF interactions in the TF layer of HuMMuS, to make a fairer comparison with state-of-the-art methods. However, the option to add links in the TF layer is provided in the released version of HuMMuS.

##### scATAC layer

scATAC data are used in this layer to infer cis-regulatory interactions using Cicero^69^. Cicero provides co-accessibility scores between peaks within given windows of the genome. We used 500kb as genomic window size for both human and mouse data, as done in^10, 69^. In addition, Cicero requires to define pseudocells, by averaging groups of N cells. In the following we used N=50, corresponding to the default Cicero value, with the only exception of the Liu dataset, where too few cells were present, thus requiring N=10. We then filtered the obtained network based on the co-accessibility scores: correlation threshold of zero for all datasets except the last dataset composed of three omics, where 0.2 is used. The obtained network is undirected and weighted.

##### scRNA layer

There are many methods to infer gene networks from scRNA data. Though it would be possible to use any network connecting genes without specifically regulatory hypotheses, we here chose to use GENIE3^70^. GENIE3 is indeed one of the most popular methods to infer GRNs from RNA and scRNA data and it was shown to have better performances than other state-of-the-art tools in^15, 16^. Being the GENIE3 network a complete one, we filtered it keeping only the 10K links with the highest weight. Of note, the network obtained by GENIE3 is here considered as an undirected and weighted network thus allowing a random walk to move from a gene to all other genes co-regulated by a common TF.

##### TF-peak bipartite

To associate TFs to potential binding regions we used the function *AddMotifs* from the Signac package^71^ and based on motifmatchr^72^. This function can be however substituted by the users with others, if needed. TF binding-motifs were obtained from JASPAR and cisBP databases^18, 68^. JASPAR motifs were obtained through the JASPAR2020 R package^73^. cisBP motifs already reformatted and deduplicated were accessed through chromVARmotifs R package^74^. To find overlap between TF binding motifs and scATAC-seq peak coordinates, elements were mapped on the genomic sequences from *BSgenome.Hsapiens.UCSC.hg38* and *BSgenome.Mmusculus.UCSC.mm10* for human and mouse, respectively. The obtained network is unweighted.

##### Peak-genes bipartite

We finally linked peaks to genes based on the distance of the peak from the transcription starting site (TSS) of the gene. We considered 500 bp before and after the TSS. The reason for the choice of this small window is due to the fact that we wanted to directly link a gene to potential promoters and leave the scATAC layer to give information on more distal regulatory regions, such as enhancers. The obtained network is unweighted.

After the reconstruction of the Heterogeneous Multilayer Network (HMLN) random walk with restart has been used for mining its information.

#### Random walk with restart (RWR)

Random walk with restart (RWR) is a stochastic process consisting in a succession of steps from one node (i.e. the seed) to a neighboring one through the network’s edges, with a probability to start again from the seed at each step. RWR can be used to explore HMLNs and to provide a measure of nodes’ closeness across the layers, ensuring the existence of a unique stationary distribution^19, 75^. To run the RWR we here used MultiXrank, a python package proposing optimized RWR on universal multilayer networks^20^.

The main parameters to run a RWR in MultiXrank are: the probability to restart from the seed and the probability to jump from one layer to another. The restart probability was set at 0.7 for all the results here presented, being this the default value in MultiXrank and also used in other RWR applications^20, 76, 77^. Concerning the probability to jump from one layer to another, we set it to be equiprobable in all layers, including the starting one. This choice is aimed at having each omic contributing equally to the results. Of note, in the HuMMuS package, when possible, we parallelized RWRs to benefit from multi-core usage.

#### Possible outputs of HuMMuS

The final outputs of HuMMuS are: (i) the prediction of the targets of a Transcription Factor (TF), based on RWRs starting from each TF in the TF layer and exploring the full network until the scRNA layer; (ii) the prediction of the peaks bound by a given TF, based on RWRs starting from each TF in the TF layer and exploring the scATAC layer; (iii) the prediction of the regulatory regions (proximal and distal enhancers) associated to a given gene, based on RWRs starting in each gene of the scRNA layer and exploring the scATAC layer; (iv) the reconstruction of Gene Regulatory Networks (GRNs), based on RWRs starting in each gene of the scRNA layer and exploring the full network until the TF layer; (v) the extraction of communities in the

GRN, reflecting tightly connected macromolecules in the HMLN frequently involved in the regulation of the same biological process or pathway^21^.

### Benchmarking settings

#### Datasets and preprocessing

The benchmarking was realized on four datasets: Chen, Liu, Duren and Semrau (see Supp Table 1). The Chen and Liu datasets consisted of paired single-cell RNA sequencing (scRNA-seq) and single-cell chromatin accessibility profiling (scATAC-seq) data from human embryonic stem cells (hESCs). Duren and Semrau consisted of unpaired scRNA-seq data from mouse embryonic stem cells (mESCs). The Semrau dataset contained only scRNA-seq data, we thus used it together with the Duren’s scATAC-seq data. Description of the data and download links can be found in Supp Table 1. Regarding data preprocessing, for both scRNA-seq and scATAC-seq data, we filtered out the features expressed in less than 1% of the cells. Gene counts were then log2-transformed and peak accessibilities were binarized by replacing the non-null values by 1.

#### Running the state-of-the-art methods

##### Pando^11^

First, unpaired datasets were computationally paired with SCOTv2^78^, running SCOTv2.align with default parameters (k=50, e=1e-3, balanced=True, rho=5e-2, normalize=True). Following the default Pando pipeline, pseudocells were then aggregated as described in https://github.com/quadbiolab/ *organoid_regulomes/blob/main/pando/pseudocells.R* to reduce data sparsity.^11^ Motifs were obtained from JASPAR2020 and cisBP, and matched to ATAC peaks with *find_motifs()*. The GRN network was finally inferred with *infer_grn()* using the parameters suggested in the Pando vignette, plus upstream = 100k, downstream = 100k and only_tss = TRUE to consider regulatory regions both downstream and upstream than the TSS, as done by the other tools here considered.

##### CellOracle^10^

We applied CellOracle as described in https://github.com/morris-lab/CellOracle. ScATAC-seq datasets were analyzed with Cicero to find co-accessible regions (co-accessibility score >0.8) in a genomic window of 500kb. Peaks co-accessible with promoters were associated with genes through CellOracle *integrate_tss_peak_with_cicero* function. Peaks were also scanned with the CellOracle *TFinfo* function and its default parameters and default motifs to identify TF binding sites, to produce TF-gene edges. Finally, the TF-gene edges were inferred by *get_links* function with alpha = 10.

##### GENIE3^70^

The R implementation of GENIE3 has been considered here. For both human and mouse datasets, we used the TFs having a known motif in JASPAR2020 or cisBP and expressed in the scRNA-seq data.

#### TF targets predictions

The aim of this first benchmark is to test the ability of different methods to predict the targets of a Transcription Factor (TF). To do this prediction with HuMMuS, we set the TFs of interest as seeds of the RWR and explored the entire HMLN until the scRNA layer to find their target genes. The probabilities of the RWR have been set as follows: (i) from the TF layer the only option was to move to the scATAC layer (as we have no link in the TF layer). We thus set a probability of 1 in the RWR to move from the TF layer to the scATAC layer; (ii) from the scATAC layer, we could stay on the layer or move either in the TF layer, either in the scRNA layer, we thus set the RWR probability to 1/3 to make all omics have the same relevance; (iii) from the scRNA layer, we could stay on the layer or move up into the scATAC layer we thus set the RWR probability to 1/2 to make all omics have the same relevance. The probability of restart was set to 0.7, default MultiXrank value. After RWR, we obtained, for each TF a ranking of putative target genes. The other state-of-the-art methods (CellOracle, GENIE3, Pando) provide a GRN, also corresponding to a list of TF-gene links reflecting a ranking of putative targets per TF. We thus evaluate performances comparing such rankings with ground-truth TF targets from^17^ that are expressed in the scRNA data. The ground truth in^17^ is composed of TF-target gene pairs for both hESCs and mESC obtained from the intersection of ChIP-seq data and perturbation experiments (impact of TFs KO/KD on gene expression).

For each method (HuMMuS, CellOracle, GENIE3, Pando) and each TF in the ground-truth, we computed Fisher tests and intersection sizes between the N top target genes and the ground-truth targets, with N varying in (3, 5, 10, 15, 20, 30, 40, 50, 75, 100). For each method, only TFs having at least 100 targets are considered. Finally, intersection performances are averaged across TFs, as TFs can vary from one method to another.

#### Regulatory regions identification

##### Predicting the peaks bounded by a TF

To predict the peaks bounded by each TF with HuMMuS, we focused on the TF layer and scATAC layer. RWRs were performed from each TF to explore the scATAC layer and find the peaks most close to them according to the RWR. The RWR probabilities were thus set to 1 for going from the TF layer to the scATAC layer (same argument for this as above); ½ to stay in the scATAC layer or move to the scRNA layer and 1 to go from the scRNA layer to the scATAC layer. The scRNA links are thus not used and the only scope of the scRNA layer is here to connect peaks associated to the regulation of the same gene. Once obtained a ranking of peaks for each TF, since the output of HuMMuS is a scoring of peaks and not a binary classification, we thresholded the ranking to only keep the top 100%, 80%, 60% or 20% of the ranking as our predictions. We then obtained Pando’s TF-peak links from the GRN post regression. Regarding CellOracle instead, TF-peak links were extracted from the backbone network, since it aggregates the peaks to calculate the TF-gene links. Since the backbone network of CellOracle is weighted according to Cicero, we further considered different Cicero thresholds (0.05, 0.2, 0.8). This list includes the default threshold of 0.8, plus additional lower thresholds since very few connections were kept with the default one. To then evaluate the quality of the obtained predictions, a ground-truth was defined from ReMap2022^22^. We thus downloaded the list of the non-redundant peaks bound per TF computed in ReMap2022, using the 37 and 193 experiments available respectively from hESCs and mESCs. Only ReMap2022 peaks overlapping with the peaks of the scATAC data were considered as part of the ground-truth. Finally, we use F1 scores to compare the peaks rankings obtained from the Pando, CellOracle and HuMMuS networks and the ground-truth peaks obtained from ReMap2022.

##### Predicting the regulatory regions (proximal and distal enhancers) associated to a gene

To predict the regulatory regions associated with a gene in HuMMuS, a RWR was computed starting from the gene as seed. No scRNA link was used, leading to a probability of 1 to go directly to the scATAC layer. Once reaching the scATAC layer, if no restart, the RWR remains in the scATAC layer with probability 1. This solution allows to explore the peaks associated with a gene based on the scATAC layer and thus potentially regulating the gene. Pando was not considered in this part of the benchmark, since it doesn’t infer peak-gene links independently from TF binding. In CellOracle, peak-gene links were extracted from the backbone networks. As for TF-regions, we considered different Cicero thresholds: 0.05, 0.2 and 0.8, with 0.8 being the default value. The obtained predictions were then compared with a ground-truth based on a combination of six enhancer databases. We first defined a list of potential enhancer-genes interactions from the union of PEGASUS^24, 27^, ENdb^23^ and EnhancerAtlas2.0^25^. We then filtered this list, keeping only the links whose enhancers were present in the union of Fantom5^28^, VISTA^29^, SCREEN.ENCODE^26^ databases. Finally, we only kept in the ground-truth enhancers overlapping with the peaks of the scATAC data. The quality of the overlap between predicted regulatory regions and the databases was finally assessed using F1 scores.

#### Community detection

As community detection methods well-suited for biological HMLG do not exist at the moment, we here compared community detection on the GRN output of HuMMuS vs. the GRNs obtained by the other methods. To obtain a GRN from HuMMuS we run, for each gene, a RWR starting from the gene as seed and arriving up to the TF layer to make TFs compete to regulate it. We thus set the probabilities to ½ to stay in the scRNA layer or to jump from it to the scATAC layer, 1/3 to jump to any of the layers from the ATAC one, and a probability of 1 to reach the scATAC layer once reaching the TF layer. Once obtained a GRN also for HuMMuS, we performed community detection on the GRNs of all methods (HuMMus, Pando, CellOracle and GENIE3). Only absolute weights were considered, all networks were filtered to the same density and community detection was finally realized with the Louvain clustering method^32^ from the networkX implementation. To find the optimal clustering resolution for each of the methods, we tested 21 values between 0 to 2 with a step size of 0.1 (see Supp Table 4). Only resolutions providing at least 10 communities out of thousands of nodes (see Supp Table 3 for details on the number of nodes per method and dataset) were considered for the following part of the analysis. We considered five different databases to evaluate the quality of the clustering : GO Cellular Component, GO Biological Process, GO Molecular Function, KEGG 2021 (human) / 2019 (mouse) and Reactome 2016^33–37^. For each method and resolution, we then used the enrichR package^79^ to find enriched pathways in each of their communities. We then counted the number and the proportion of communities significantly enriched (p-value < 0.05 in the results presented Fig. 4) in at least one geneset of the database. For each method, we selected the resolution returning best performances.

### HuMMuS applied to mouse cortex profiled for scRNA, scATAC and scnmC

#### HuMMuS application from HMLN reconstruction to GRN extraction

To illustrate the potential of HuMMuS we used a single-cell dataset of cortical neurons composed of snmC, snATAC-seq and scRNA-seq. The data were downloaded from^38–40^. The snmC dataset was composed of 46,714 genes and 3386 cells; scRNA-seq was composed of 25,299 genes and 55,803 cells and scATAC-seq was composed of 155,093 peaks and 2317 cells. For scATAC and scRNA, we used preprocessed data in the h5ad files accessible at https://scglue.readthedocs.io/en/latest/data.html under the names *Saunders-2018* and *10x-Multiome-Pbmc10k*, while for snmC data, we used mCH methylation averaged per gene body (gene_level_mouse.txt) available at https://brainome.ucsd.edu/annoj/brain_single_nuclei/snmcSeq_processed_data.tar.g <u>z</u> and retained only the features expressed in more than 3% of the cells.

We then used HuMMuS to contract a HMLN consisting of four layers: a TF layer, a snmC layer, a scATAC layer, and a scRNA layer. To follow transcriptional regulation structure we placed the scmC layer in the middle between the scATAC layer and the scRNA layer. We didn’t link the snmC layer to the TF layer because TF binding motifs are specific to small regions, making gene bodies too large for precise binding motifs. As in the benchmark, we didn’t put links in the TF layer. For the scATAC layer we used Cicero setting a co-accessibility score threshold at 0.2, as almost all correlations were above 0. The scRNA layer was computed with the python version of GRNBoost2, GENIE3 did not manage to get results on such a big dataset. Then the 50k links with the highest weights were kept. For the scmC layer, since we did not find methods designed to infer networks on methylation data, we used partial correlation from the pingouin0.5.3 python package, accessible at https://github.com/raphaelvallat/pingouin/tree/master. All the links with an absolute corrected correlation above 0.3 were kept. The inter-layer connections not involving the snmC layer were structured as in the benchmark. The connections between the snmC layer and the scATAC layer were set based on the distance of the scATAC peaks from the transcription start site (TSS) of the genes, nodes of the snmC layer (500 bp before and after the TSS). The connections between the snmC layer and the scRNA layer were just based on gene-gene correspondence.

After HMLG construction, using RWR from the gene layer up to the TF layer, we reconstructed a GRN. To give the same importance to each modality, the probability to go to any possible layer was the same. For the scATAC layer, we then have a probability of ¼ to go to each of the other layers or to stay in. For the scRNA layer and the snmC layer, we have a probability of ⅓ to stay in the layer, to move to the atac-layer or to move to the other gene-node network. Finally, from the TFs layer it’s only possible to jump to the atac-layer.

#### Data analysis with the obtained GRN

Starting from the GRN provided by HuMMuS, we isolated regulons, corresponding to TFs and their linked genes, and evaluated their activity in scRNA data using the unilinear model implemented in Decoupler^41^. UMAP was then run on such an activity matrix to test the ability of the obtained regulons to cluster cells according to their cortical neuron sub-population of origin. Finally, TF activities were used to find top marker regulons of each cortical neuron sub-population focusing on the top 10 regulons per cortical sub-population.

## Acknowledgements

This work was supported by funding from the Agence Nationale de la Recherche (ANR) JCJC project scMOmix and the French government under management of Agence Nationale de la Recherche as part of the ‘Investissements d’avenir’ program, reference ANR19-P3IA-0001 (PRAIRIE 3IA Institute).

## Author contributions

L.C. and R. T. designed and planned the study. L.C. wrote the paper. R.T. developed the tool and performed most of the analyses. IM.D. contributed to the analyses.

## Disclosure and competing interests statement

The authors declare no competing interests.

## Data Availability

The code to run HuMMuS is available at https://github.com/cantinilab/HuMMuS together with tutorials. For the input data all details to access them are reported in the second column of Supp Table 1 plus links to access the preprocessed data are available at https://github.com/cantinilab/HuMMuS.

## Notes

https://github.com/cantinilab/HuMMuS

### Competing Interest Statement

The authors have declared no competing interest.

