## Supplementary Material for "Molecular mechanisms reconstruction from single-cell multi-omics data with HuMMuS"

### Supplementary figures

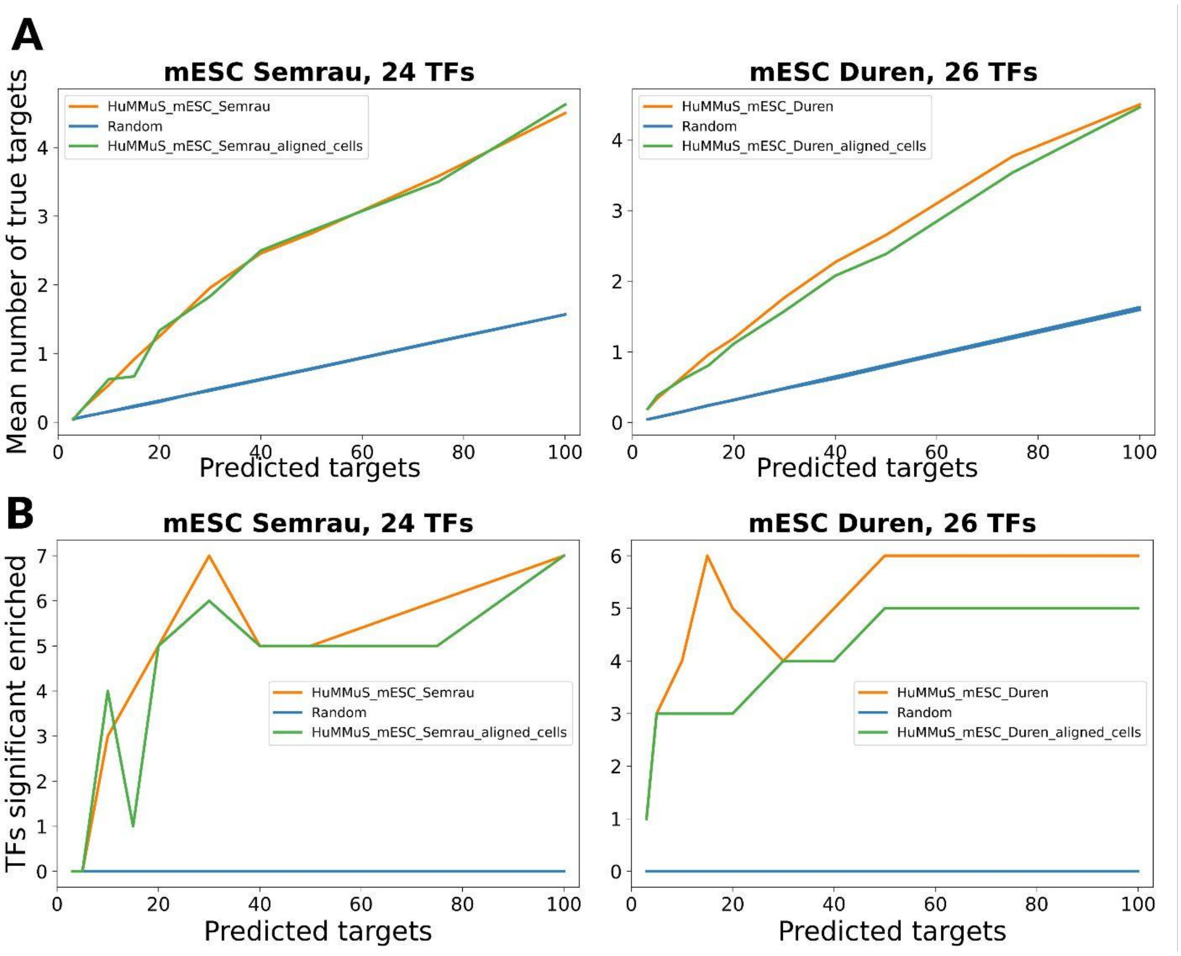


**Supp Figure 1 - Transcription Factor (TF) targets prediction with and without cell pairing** (A) average number of correctly predicted targets per TF. (B) number of TFs having a significant amount of correctly predicted targets (Fisher's test p-value <0.05). In (A-B) different colors correspond to different methods: orange (HuMMus on unpaired scATAC and scRNA-seq data), green (HuMMuS on paired scATAC and scRNA-seq data), blue (Random)

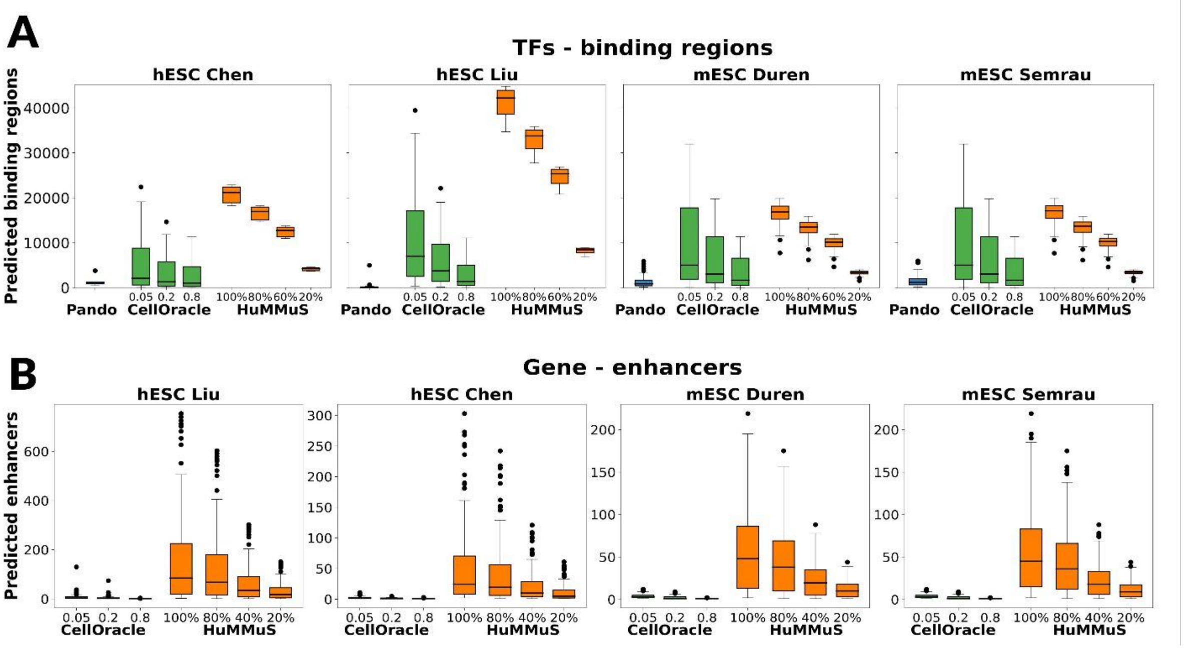


**Supp Figure 2 - Binding regions and regulatory regions prediction with and without cell pairing.** F1 score distributions of the intersection between the ground-truth of TF-peak associations and those inferred by HuMMuS. Different colors correspond to different data processing: orange (HuMMuS on unpaired scATAC and scRNAseq data), blue (HuMMuS on paired scATAC and scRNA-seq data).

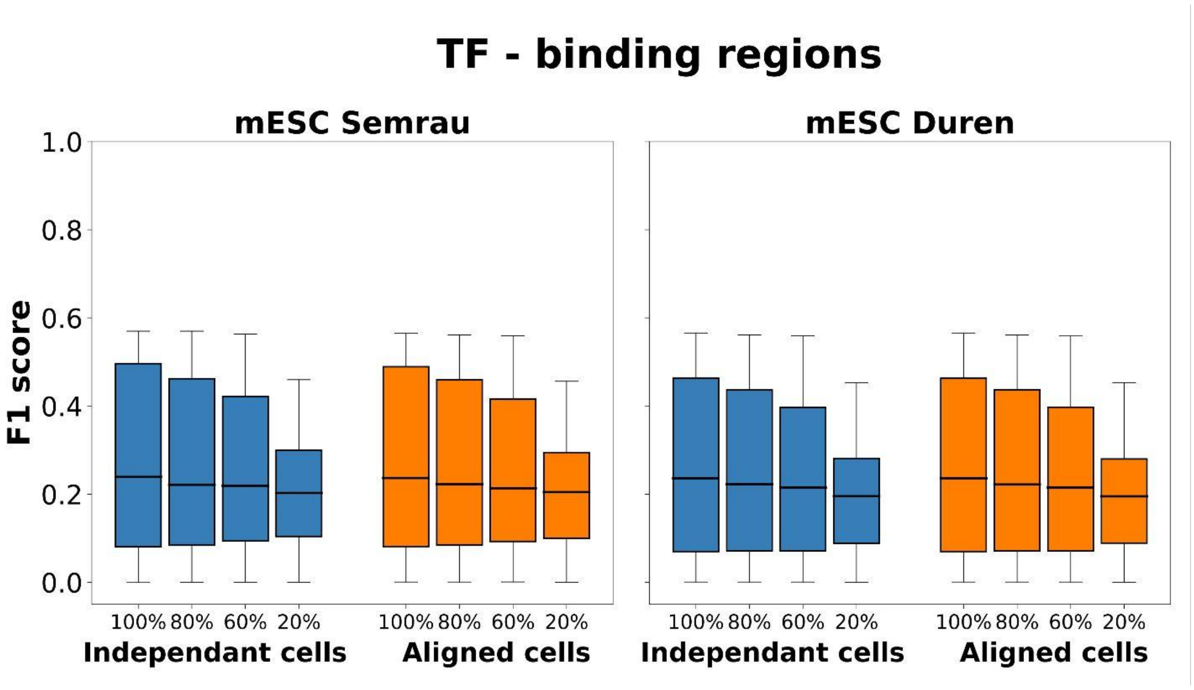


**Supp Figure 3 - number of regions detected per TF-regions and regions - peaks methods** (A) Distribution of the number of binding regions per tf inferred by Pando, CellOracle and HuMMuS; (B) Distribution of the number of regulatory regions per gene inferred by CellOracle and HuMMuS. In (A-B) different colors correspond to different methods: orange (HuMMuS), blue (Pando), green (CellOracle).

**Supplementary Table 1. Collection of publicly data used in this study.**(A) Description of the four datasets to benchmark HuMMuS in respect to state-of-the-art methods. It notably contains accessibility identifier/link and related originial publication. (B) List of the ground truth used to benchmark targets genes of transcription factors for the mESC and hESC datasets described above. (C) Databases of enhancers and regulatory regions combined to evaluate methods to retrieve regulatory regions important for each genes. (D) List of the databases used to evaluate enrichment of community detected from GRN benchmarking. (E) List and accessibility links to the data used for the 3 omics multilayer reconstruction. We used the table already preprocessed and formated by Cao et al., 2022.

**A. Benchmark data: Gene expression and chromatin accessibility datasets**

| Dataset name | Data accession | Associated publication |
|---|---|---|
| hESC_Chen | NCBI (GSE126074): | (Chen et al., 2019) |
| scRNA | 8595 features and 385 cells | Obtained by SNARE-seq. cell line mixture SNAREseq cDNA counts and chromatin counts, cell labels for filtering (ftp://ftp.ebi.ac.uk/ pub/ databases/mofa/snare_seq/cell_metadata.txt). Here, we only use H1 cells. |
| scATAC | 36954 features and 385 cells | |
| hESC_Liu | [https://github.com/hdsu-bioquant/scCAT/blob/master/data/HumanEmbryo/](https://github.com/hdsu-bioquant/scCAT/blob/master/data/HumanEmbryo/). | (Liu et al., 2019) |
| scRNA | 23153 features and 72 cells | Obtained by scCAT. Human embryo RNA-seq counts, ATAC-seq counts, and annotation data downloaded from github link |
| scATAC | 68952 features and 72 cells | |
| mESC_Duren | NCBI (GSE115968): scRNA-seq_RA_D4, and NCBI (GSE115970): scATAC-seq_RA_D4 | (Duren et al., 2018; Zeng et al., 2019) |
| scRNA | 15299 features and 464 cells | 415 scATAC-seq samples generated for the retinoic acid-induced mESC differenation at day 4. |
| scATAC | 23176 features and 414 cells | 464 scRNA-seq samples generated for the retinoic acid-induced mESC differenation at day 4. |
| mESC_Semrau | NCBI (GSM2098553) part of (GSE79578): scrbseq_96h (scRNA-seq), but no scATAC-seq. We use the scATAC-seq from mESC_Duren, NCBI (GSE115970). | (Semrau et al., 2017) |
| scRNA | 10243 features and 384 cells | scRNA seq data generated alone. scATAC-seq data of Semrau et al. has been used as peak layer |

**B. Ground truth table of target genes of transcription factors**

| Dataset name | Data accession link | Associated publication |
|---|---|---|
| mESC & hESC TF GT | [https://zenodo.org/record/5909090/files/gold_standard_datasets.zip?download=1](https://zenodo.org/record/5909090/files/gold_standard_datasets.zip?download=1), Tables used hESC_chipunion_KDUnion_intersect.txt, and mESC_chipunion_KDUnion_intersect.txt, downloaded: 6th April 2022 | (Stone et al., 2022) |

**C. Databases used to construct Ground truth table of enhancers for genes**

| Databases name | Data accession | Associated publication |
|---|---|---|
| EnhancerAtlas2.0 | [http://www.enhanceratlas.org/indexv2.php](http://www.enhanceratlas.org/indexv2.php), Table used (ESC_neuron_EP.txt, ESC_Bruce4_EP.txt, ESC_J1_EP.txt, and ESC_KH2_EP.txt): enhancer-gene interactions - Homo sapiens (hg19: ESC_neuron) and Mus musculus (mm9: ESC_Bruce4, ESC_J1, ESC_KH2), Downloaded: 4th April 2022, Preprocessing: table formatting, convert hg19 to hg38, mm9 to mm10, and ENSEMBL Gene IDs to Gene Symbol | (Gao & Qian, 2020) |
| ENdb | [http://www.licpathway.net/ENdb/](http://www.licpathway.net/ENdb/), Table used (ENdb_enhancer.txt): All the experimentally confirmed enhancers (hg19 and mm10), Downloaded: 4th April 2022, Preprocessing: convert hg19 to hg38 | (Bai et al., 2020) |
| SCREEN (ENCODE) | [http://screen.encodeproject.org](http://screen.encodeproject.org), Table used (GRCh38-cCREs.bed.html and mm10-ccREs.bed.html): all human cCREs (hg38) and all mouse cCREs (mm10), Downloaded: 4th April 2022, Preprocessing: none | (The ENCODE Project Consortium et al., 2020) |
| VISTA | [http://enhancer.lbl.gov](http://enhancer.lbl.gov), Table used (imagedb3.pl.html): all 3281 elements (hg19 and mm9), Downloaded: 4th April 2022, Preprocessing: delete sequences, bring into table form, convert hg19 to hg38, mm9 to mm10 | (Visel et al., 2007) |
| PEGASUS | [ftp://ftp.biologie.ens.fr/pub/dyogen/PEGASUS/](ftp://ftp.biologie.ens.fr/pub/dyogen/PEGASUS/), Table used (hg19_CNEs_PEGASUS.data.gz): PEGASUS predictions for the human genome (hg19), Downloaded: 4th April 2022, Preprocessing: convert hg19 to hg38, hg19 to mm10, and ENSEMBL Gene IDs to Gene Symbol | (Naville et al., 2015; Clément et al., 2020) |
| Fantom5 | [https://slidebase.binf.ku.dk/human_enhancers/presets](https://slidebase.binf.ku.dk/human_enhancers/presets), Table used (hg19_enhancer_promoter_correlations_distances_cell_type.txt.gz): Enhancer-Promoter Cell Type Associations in 5.Enhancer - FANTOM Robust Promoter associations, Downloaded: 12th May 2022, Preprocessing: convert hg19 to hg38, hg19 to mm10 for enhancer and also promoter regions | (Forrest et al., 2014) |

**D. Databases for gene enrichment analyses included in enrichR (Chen et al., 2013; Kuleshov et al., 2016; Xie et al., 2021)**

| Databases name | Data accession | Associated publication |
|---|---|---|
| Gene Ontology | GO_Biological_Processes_2021, GO_Cellular_Component_2021, and GO_Molecular_Function_2021 | (Ashburner et al., 2000; Gene Ontology Consortium, 2021) |
| Kyoto Encyclopedia of Genes and | KEGG_2021_Human and KEGG_2019_Mouse | (Kanehisa & Goto, 2000; Kanehisa, 2019; Kanehisa et al., 2021) |
| Reactome | Reactome_2016 (citation refers to latest Reactome version not used within enrichR) | (Gillespie et al., 2022) |

**E. Dataset to test inclusion of 4th layer: Gene expression, chromatin accessibility, and HiC data**

| Dataset name | Data accession | Associated publication |
|---|---|---|
| scRNA mouse cortex | [http://download.gao-lab.org/GLUE/dataset/Saunders-2018.h5ad](http://download.gao-lab.org/GLUE/dataset/Saunders-2018.h5ad) | Saunders et al., 2018 |
| scATAC mouse cortex | [http://download.gao-lab.org/GLUE/dataset/10x-ATAC-Brain5k.h5ad](http://download.gao-lab.org/GLUE/dataset/10x-ATAC-Brain5k.h5ad) | [https://support.10xgenomics.com/single-cell-atac/datasets/1.1.0/atac_v1_adult_brain_fresh_5k](https://support.10xgenomics.com/single-cell-atac/datasets/1.1.0/atac_v1_adult_brain_fresh_5k) |
| snmC mouse cortex | [http://download.gao-lab.org/GLUE/dataset/Luo-2017.h5ad](http://download.gao-lab.org/GLUE/dataset/Luo-2017.h5ad) | Luo et al., 2017 |

**Supplementary Table 2. General description of the different multilayers components.** This table contains number of nodes and edges in each component of the multilayers analysed in this article

| | Layer / Bipartite | 3 omics mouse cortex | hESC Chen | hESC Liu | mESC Duren | mESC Semrau |
|---|---|---|---|---|---|---|
| **TF layer** | Number of nodes | 717 | 220 | 670 | 607 | 334 |
| **scATAC layer** | Number of nodes | 82108 | 25102 | 48207 | 20779 | 20779 |
| | Number of edges | 302651 | 96104 | 439628 | 72986 | 72986 |
| **scRNA layer** | Number of nodes | 9349 | 5095 | 5712 | 4520 | 5695 |
| | Number of edges | 50000 | 10000 | 10000 | 10000 | 10000 |
| **snmC layer** | Number of nodes | 5494 | | | | |
| | Number of edges | 558845 | | | | |
| **TF --> scATAC bipartite** | Number of sources | 717 | 220 | 670 | 607 | 334 |
| | Number of targets | 153507 | 25102 | 48207 | 20779 | 20779 |
| | Number of edges | 10685078 | 619902 | 3520936 | 1373435 | 792771 |
| **scATAC--> scRNA bipartite** | Number of sources | 36494 | 4989 | 3624 | 3036 | 4772 |
| | Number of targets | 16214 | 3780 | 2086 | 2661 | 4367 |
| | Number of edges | 36494 | 5230 | 3690 | 3096 | 4986 |
| **scATAC --> snmC bipartite** | Number of sources | 35210 | | | | |
| | Number of targets | 17284 | | | | |
| | Number of edges | 38299 | | | | |
| **snmC --> scRNA bipartite** | Number of sources | 24504 | | | | |
| | Number of targets | 24504 | | | | |
| | Number of edges | 24504 | | | | |

**Supplementary Table 3. Density comparison between the GRNs of the different methods.** Table containing the number of TFs, genes, edges and the density of the GRNs defined by each method for the benchmark.

| | | Number of TFs | Number of genes | Number of edges | Density |
|---|---|---|---|---|---|
| **hESC_Chen** | **celloracle** | 252 | 5925 | 369196 | 0.24731 |
| | **HuMMuS** | 220 | 5095 | 1120680 | 1 |
| | **pando** | 220 | 8152 | 270817 | 0.151023 |
| | **genie3** | 220 | 8595 | 1875712 | 0.992083 |
| **hESC_Liu** | **hmln** | 670 | 5733 | 3795184 | 0.988216 |
| | **HuMMuS** | 670 | 5733 | 3801638 | 0.989896 |
| | **pando** | 517 | 15517 | 232767 | 0.029017 |
| | **genie3** | 670 | 23153 | 12424093 | 0.800943 |
| **mESC_Semrau** | **celloracle** | 388 | 7662 | 706606 | 0.237717 |
| | **HuMMuS** | 334 | 5695 | 1901796 | 1 |
| | **pando** | 334 | 9658 | 695894 | 0.215752 |
| | **genie3** | 334 | 10243 | 3300296 | 0.964765 |
| **mESC_Duren** | **celloracle** | 664 | 10473 | 1337231 | 0.192313 |
| | **HuMMuS** | 607 | 4570 | 2741871 | 0.988638 |
| | **pando** | 602 | 14045 | 1240683 | 0.146748 |
| | **genie3** | 607 | 15299 | 8777184 | 0.945218 |

**Supplementary Table 4.** Number of significantly enriched communities (p value < 0.05) detected for each methods, resolution, and databases used.

| | | | Resolution | | | | | | | | | | | | | | | | | | | | |
|---|---|---|---|---|---|---|---|---|---|---|---|---|---|---|---|---|---|---|---|---|---|---|---|
| | | | 0 | 0.1 | 0.2 | 0.3 | 0.4 | 0.5 | 0.6 | 0.7 | 0.8 | 0.9 | 1 | 1.1 | 1.2 | 1.3 | 1.4 | 1.5 | 1.6 | 1.7 | 1.8 | 1.9 | 2 |
| **hESC_Chen** | **celloracle** | GO_Biological_Process_2021 | 1 | 1 | 1 | 1 | 1 | 1 | 1 | 1 | 1 | 4 | 10 | 12 | 12 | 22 | 12 | 20 | 31 | 30 | 32 | 32 | 30 |
| | | GO_Cellular_Component_2021 | 1 | 1 | 1 | 1 | 1 | 1 | 1 | 1 | 1 | 4 | 9 | 12 | 13 | 20 | 18 | 23 | 31 | 29 | 31 | 31 | 35 |
| | | GO_Molecular_Function_2021 | 1 | 1 | 1 | 1 | 1 | 1 | 1 | 1 | 1 | 4 | 10 | 9 | 17 | 20 | 19 | 23 | 28 | 39 | 29 | 35 | 33 |
| | | KEGG_2021_Human | 1 | 1 | 1 | 1 | 1 | 1 | 1 | 1 | 1 | 4 | 5 | 3 | 7 | 15 | 6 | 10 | 10 | 12 | 14 | 7 | 8 |
| | | Reactome_2016 | 1 | 1 | 1 | 1 | 1 | 1 | 1 | 1 | 1 | 4 | 9 | 14 | 15 | 22 | 16 | 22 | 25 | 21 | 27 | 23 | 36 |
| | **pando** | GO_Biological_Process_2021 | 1 | 2 | 2 | 2 | 2 | 2 | 2 | 2 | 2 | 2 | 2 | 3 | 3 | 4 | 4 | 3 | 3 | 2 | 2 | 2 | 2 |
| | | GO_Cellular_Component_2021 | 1 | 2 | 2 | 2 | 2 | 2 | 2 | 2 | 2 | 2 | 2 | 2 | 2 | 3 | 3 | 2 | 2 | 0 | 1 | 1 | 1 |
| | | GO_Molecular_Function_2021 | 1 | 2 | 2 | 2 | 2 | 2 | 2 | 2 | 2 | 2 | 2 | 3 | 3 | 4 | 4 | 2 | 3 | 2 | 2 | 2 | 2 |
| | | KEGG_2021_Human | 1 | 2 | 2 | 2 | 2 | 2 | 2 | 2 | 2 | 2 | 2 | 1 | 2 | 4 | 3 | 2 | 2 | 0 | 1 | 1 | 1 |
| | | Reactome_2016 | 1 | 2 | 2 | 2 | 2 | 2 | 2 | 2 | 2 | 2 | 2 | 2 | 3 | 4 | 4 | 3 | 3 | 2 | 2 | 2 | 2 |
| | **genie3** | GO_Biological_Process_2021 | 1 | 1 | 1 | 1 | 1 | 1 | 1 | 1 | 1 | 4 | 10 | 8 | 5 | 8 | 13 | 19 | 20 | 13 | 21 | 22 | 24 |
| | | GO_Cellular_Component_2021 | 1 | 1 | 1 | 1 | 1 | 1 | 1 | 1 | 1 | 6 | 7 | 10 | 8 | 10 | 15 | 20 | 15 | 21 | 20 | 18 | 21 |
| | | GO_Molecular_Function_2021 | 1 | 1 | 1 | 1 | 1 | 1 | 1 | 1 | 1 | 5 | 9 | 8 | 7 | 12 | 12 | 15 | 17 | 24 | 22 | 24 | 29 |
| | | KEGG_2021_Human | 1 | 1 | 1 | 1 | 1 | 1 | 1 | 1 | 1 | 4 | 4 | 2 | 1 | 6 | 7 | 12 | 10 | 11 | 14 | 14 | 11 |
| | | Reactome_2016 | 1 | 1 | 1 | 1 | 1 | 1 | 1 | 1 | 1 | 5 | 11 | 10 | 5 | 7 | 19 | 16 | 15 | 15 | 20 | 15 | 20 |
| | **HuMMuS** | GO_Biological_Process_2021 | 1 | 1 | 1 | 1 | 1 | 1 | 1 | 1 | 2 | 4 | 5 | 3 | 6 | 8 | 10 | 21 | 16 | 19 | 18 | 24 | 32 |
| | | GO_Cellular_Component_2021 | 1 | 1 | 1 | 1 | 1 | 1 | 1 | 1 | 2 | 4 | 4 | 6 | 6 | 8 | 7 | 17 | 14 | 13 | 20 | 21 | 22 |
| | | GO_Molecular_Function_2021 | 1 | 1 | 1 | 1 | 1 | 1 | 1 | 1 | 2 | 4 | 5 | 3 | 6 | 6 | 10 | 13 | 17 | 18 | 16 | 17 | 24 |
| | | KEGG_2021_Human | 1 | 1 | 1 | 1 | 1 | 1 | 1 | 1 | 2 | 2 | 4 | 4 | 6 | 5 | 11 | 11 | 9 | 14 | 16 | 7 | 15 |
| | | Reactome_2016 | 1 | 1 | 1 | 1 | 1 | 1 | 1 | 1 | 2 | 4 | 6 | 8 | 8 | 8 | 10 | 15 | 15 | 25 | 20 | 11 | 23 |
| **hESC_Liu** | **celloracle** | GO_Biological_Process_2021 | 1 | 1 | 1 | 1 | 1 | 1 | 1 | 3 | 3 | 6 | 8 | 10 | 17 | 18 | 19 | 20 | 25 | 37 | 39 | 33 | 45 |
| | | GO_Cellular_Component_2021 | 1 | 1 | 1 | 1 | 1 | 1 | 1 | 2 | 3 | 6 | 8 | 7 | 13 | 14 | 16 | 24 | 19 | 28 | 35 | 16 | 24 |
| | | GO_Molecular_Function_2021 | 1 | 1 | 1 | 1 | 1 | 1 | 1 | 2 | 3 | 6 | 8 | 9 | 13 | 17 | 19 | 22 | 26 | 23 | 40 | 29 | 36 |
| | | KEGG_2021_Human | 1 | 1 | 1 | 1 | 1 | 1 | 1 | 2 | 3 | 6 | 6 | 8 | 10 | 8 | 8 | 14 | 15 | 13 | 15 | 21 | 18 |
| | | Reactome_2016 | 1 | 1 | 1 | 1 | 1 | 1 | 1 | 2 | 3 | 3 | 4 | 9 | 10 | 9 | 16 | 18 | 19 | 30 | 33 | 19 | 30 |
| | **pando** | GO_Biological_Process_2021 | 1 | 1 | 3 | 4 | 5 | 4 | 2 | 3 | 2 | 3 | 1 | 2 | 2 | 4 | 4 | 3 | 1 | 3 | 4 | 2 | 1 |
| | | GO_Cellular_Component_2021 | 1 | 1 | 3 | 5 | 5 | 4 | 0 | 1 | 1 | 1 | 1 | 1 | 4 | 3 | 3 | 3 | 3 | 2 | 3 | 2 | 2 |
| | | GO_Molecular_Function_2021 | 1 | 1 | 3 | 4 | 5 | 4 | 0 | 3 | 2 | 2 | 1 | 2 | 2 | 3 | 6 | 0 | 2 | 1 | 4 | 2 | 1 |
| | | KEGG_2021_Human | 1 | 1 | 3 | 4 | 5 | 4 | 0 | 1 | 1 | 2 | 1 | 2 | 3 | 2 | 4 | 2 | 1 | 1 | 3 | 2 | 2 |
| | | Reactome_2016 | 1 | 1 | 3 | 4 | 5 | 4 | 3 | 4 | 3 | 1 | 1 | 2 | 4 | 3 | 5 | 3 | 2 | 5 | 3 | 3 | 4 |
| | **genie3** | GO_Biological_Process_2021 | 0 | 0 | 0 | 0 | 1 | 1 | 1 | 1 | 0 | 0 | 0 | 0 | 2 | 6 | 3 | 2 | 3 | 3 | 2 | 6 | 5 |
| | | GO_Cellular_Component_2021 | 0 | 0 | 0 | 0 | 1 | 1 | 1 | 1 | 0 | 0 | 0 | 0 | 2 | 5 | 2 | 1 | 2 | 3 | 3 | 6 | 4 |
| | | GO_Molecular_Function_2021 | 0 | 0 | 0 | 0 | 1 | 1 | 1 | 1 | 0 | 0 | 0 | 0 | 2 | 5 | 2 | 1 | 2 | 3 | 2 | 7 | 5 |
| | | KEGG_2021_Human | 0 | 0 | 0 | 0 | 1 | 1 | 1 | 1 | 0 | 0 | 0 | 1 | 2 | 7 | 4 | 1 | 2 | 4 | 3 | 7 | 4 |
| | | Reactome_2016 | 0 | 0 | 0 | 0 | 1 | 1 | 1 | 1 | 0 | 0 | 0 | 0 | 3 | 5 | 2 | 2 | 3 | 3 | 3 | 6 | 4 |
| | **HuMMuS** | GO_Biological_Process_2021 | 1 | 1 | 1 | 1 | 1 | 1 | 1 | 1 | 2 | 6 | 8 | 10 | 16 | 20 | 21 | 28 | 31 | 32 | 46 | 41 | 51 |
| | | GO_Molecular_Function_2021 | 1 | 1 | 1 | 1 | 1 | 1 | 1 | 1 | 2 | 6 | 8 | 10 | 16 | 19 | 23 | 28 | 33 | 36 | 41 | 50 | 48 |
| | | KEGG_2021_Human | 1 | 1 | 1 | 1 | 1 | 1 | 1 | 1 | 2 | 5 | 8 | 8 | 12 | 12 | 10 | 16 | 13 | 18 | 21 | 17 | 21 |
| | | Reactome_2016 | 1 | 1 | 1 | 1 | 1 | 1 | 1 | 1 | 2 | 5 | 6 | 8 | 14 | 16 | 14 | 20 | 22 | 22 | 26 | 35 | 28 |
| | | GO_Cellular_Component_2021 | 0 | 0 | 0 | 0 | 1 | 1 | 1 | 1 | 2 | 6 | 8 | 8 | 11 | 11 | 13 | 16 | 15 | 19 | 17 | 15 | 18 |
| **mESC_Duren** | **celloracle** | GO_Biological_Process_2021 | 1 | 1 | 1 | 1 | 1 | 1 | 1 | 1 | 1 | 3 | 8 | 11 | 13 | 22 | 31 | 39 | 59 | 54 | 62 | 61 | 58 |
| | | GO_Cellular_Component_2021 | 1 | 1 | 1 | 1 | 1 | 1 | 1 | 1 | 1 | 3 | 8 | 11 | 10 | 18 | 24 | 36 | 32 | 39 | 45 | 36 | 35 |
| | | GO_Molecular_Function_2021 | 1 | 1 | 1 | 1 | 1 | 1 | 1 | 1 | 1 | 3 | 8 | 6 | 15 | 22 | 29 | 31 | 33 | 49 | 49 | 43 | 53 |
| | | KEGG_2019_Mouse | 1 | 1 | 1 | 1 | 1 | 1 | 1 | 1 | 1 | 3 | 8 | 10 | 13 | 17 | 26 | 29 | 25 | 36 | 26 | 30 | 29 |
| | | Reactome_2016 | 1 | 1 | 1 | 1 | 1 | 1 | 1 | 1 | 1 | 3 | 8 | 13 | 16 | 25 | 31 | 42 | 35 | 43 | 50 | 49 | 51 |
| | **pando** | GO_Biological_Process_2021 | 1 | 1 | 2 | 1 | 2 | 2 | 2 | 2 | 2 | 2 | 2 | 9 | 9 | 15 | 17 | 10 | 14 | 13 | 12 | 14 | 14 |
| | | GO_Cellular_Component_2021 | 1 | 1 | 2 | 1 | 1 | 2 | 2 | 2 | 2 | 2 | 2 | 3 | 4 | 6 | 7 | 6 | 6 | 8 | 5 | 11 | 4 |
| | | GO_Molecular_Function_2021 | 1 | 1 | 2 | 2 | 2 | 2 | 2 | 2 | 2 | 2 | 2 | 9 | 14 | 19 | 19 | 21 | 18 | 20 | 17 | 21 | 18 |
| | | KEGG_2019_Mouse | 1 | 2 | 2 | 1 | 2 | 2 | 2 | 2 | 2 | 2 | 2 | 3 | 4 | 7 | 3 | 4 | 7 | 7 | 3 | 7 | 4 |
| | | Reactome_2016 | 1 | 2 | 2 | 1 | 2 | 2 | 2 | 2 | 2 | 2 | 2 | 5 | 6 | 6 | 8 | 8 | 8 | 8 | 5 | 8 | 9 |
| | **genie3** | GO_Biological_Process_2021 | 1 | 1 | 1 | 1 | 1 | 1 | 2 | 3 | 4 | 3 | 4 | 5 | 6 | 7 | 7 | 6 | 10 | 10 | 9 | 14 | 13 |
| | | GO_Cellular_Component_2021 | 1 | 1 | 1 | 1 | 1 | 1 | 2 | 3 | 4 | 3 | 4 | 5 | 5 | 8 | 7 | 12 | 5 | 15 | 9 | 13 | 9 |
| | | GO_Molecular_Function_2021 | 1 | 1 | 1 | 1 | 1 | 1 | 2 | 3 | 4 | 3 | 4 | 5 | 6 | 8 | 8 | 5 | 11 | 10 | 13 | 17 | 15 |
| | | KEGG_2019_Mouse | 1 | 1 | 1 | 1 | 1 | 1 | 2 | 3 | 4 | 3 | 4 | 5 | 5 | 8 | 9 | 7 | 5 | 11 | 7 | 12 | 13 |
| | | Reactome_2016 | 1 | 1 | 1 | 1 | 1 | 1 | 2 | 3 | 4 | 3 | 4 | 5 | 6 | 9 | 8 | 6 | 11 | 16 | 9 | 20 | 11 |
| | **HuMMuS** | GO_Biological_Process_2021 | 1 | 1 | 1 | 1 | 1 | 1 | 1 | 1 | 2 | 3 | 8 | 14 | 16 | 23 | 22 | 30 | 31 | 32 | 41 | 42 | 45 |
| | | GO_Cellular_Component_2021 | 1 | 1 | 1 | 1 | 1 | 1 | 1 | 1 | 2 | 3 | 8 | 7 | 8 | 13 | 11 | 12 | 15 | 20 | 26 | 18 | 30 |
| | | GO_Molecular_Function_2021 | 1 | 1 | 1 | 1 | 1 | 1 | 1 | 1 | 2 | 3 | 8 | 13 | 17 | 21 | 24 | 25 | 26 | 31 | 36 | 43 | 40 |
| | | KEGG_2019_Mouse | 1 | 1 | 1 | 1 | 1 | 1 | 1 | 1 | 2 | 3 | 5 | 7 | 13 | 17 | 16 | 20 | 17 | 23 | 30 | 24 | 34 |
| | | Reactome_2016 | 1 | 1 | 1 | 1 | 1 | 1 | 1 | 1 | 2 | 3 | 8 | 13 | 15 | 21 | 19 | 24 | 18 | 29 | 34 | 37 | 39 |
| **mESC_Semrau** | **celloracle** | GO_Biological_Process_2021 | 1 | 1 | 1 | 1 | 1 | 1 | 1 | 1 | 1 | 4 | 6 | 11 | 24 | 29 | 30 | 29 | 49 | 35 | 46 | 51 | 45 |
| | | GO_Cellular_Component_2021 | 1 | 1 | 1 | 1 | 1 | 1 | 1 | 1 | 1 | 4 | 6 | 11 | 16 | 26 | 31 | 16 | 26 | 36 | 47 | 27 | 27 |
| | | GO_Molecular_Function_2021 | 1 | 1 | 1 | 1 | 1 | 1 | 1 | 1 | 1 | 4 | 5 | 8 | 13 | 19 | 28 | 29 | 33 | 38 | 43 | 41 | 43 |
| | | KEGG_2019_Mouse | 1 | 1 | 1 | 1 | 1 | 1 | 1 | 1 | 1 | 4 | 6 | 7 | 11 | 13 | 12 | 26 | 27 | 28 | 31 | 23 | 24 |
| | | Reactome_2016 | 1 | 1 | 1 | 1 | 1 | 1 | 1 | 1 | 1 | 4 | 6 | 13 | 21 | 25 | 23 | 33 | 37 | 42 | 39 | 41 | 39 |
| | **pando** | GO_Biological_Process_2021 | 1 | 1 | 2 | 2 | 2 | 2 | 2 | 2 | 2 | 2 | 2 | 8 | 12 | 18 | 11 | 15 | 14 | 18 | 16 | 15 | 21 |
| | | GO_Cellular_Component_2021 | 1 | 2 | 2 | 2 | 2 | 2 | 2 | 2 | 2 | 2 | 2 | 6 | 10 | 11 | 10 | 9 | 10 | 16 | 15 | 12 | 20 |
| | | GO_Molecular_Function_2021 | 1 | 1 | 1 | 2 | 2 | 2 | 2 | 2 | 2 | 2 | 2 | 11 | 15 | 17 | 16 | 21 | 19 | 21 | 20 | 18 | 23 |
| | | KEGG_2019_Mouse | 1 | 1 | 2 | 2 | 1 | 2 | 2 | 2 | 2 | 2 | 2 | 7 | 8 | 9 | 8 | 12 | 10 | 14 | 10 | 11 | 9 |
| | | Reactome_2016 | 1 | 2 | 2 | 2 | 2 | 2 | 2 | 2 | 2 | 2 | 2 | 8 | 10 | 14 | 9 | 13 | 15 | 14 | 13 | 14 | 12 |
| | **genie3** | GO_Biological_Process_2021 | 1 | 1 | 1 | 1 | 1 | 1 | 1 | 1 | 2 | 2 | 6 | 2 | 5 | 10 | 7 | 10 | 17 | 10 | 8 | 13 | 16 |
| | | GO_Cellular_Component_2021 | 1 | 1 | 1 | 1 | 1 | 1 | 1 | 1 | 2 | 2 | 6 | 2 | 6 | 2 | 8 | 11 | 15 | 25 | 16 | 25 | 19 |
| | | GO_Molecular_Function_2021 | 1 | 1 | 1 | 1 | 1 | 1 | 1 | 1 | 2 | 2 | 6 | 3 | 7 | 10 | 9 | 7 | 18 | 11 | 12 | 28 | 16 |
| | | KEGG_2019_Mouse | 1 | 1 | 1 | 1 | 1 | 1 | 1 | 1 | 2 | 2 | 7 | 2 | 6 | 5 | 6 | 7 | 10 | 9 | 13 | 13 | 9 |
| | | Reactome_2016 | 1 | 1 | 0 | 1 | 1 | 1 | 1 | 1 | 2 | 2 | 7 | 10 | 8 | 3 | 8 | 7 | 18 | 13 | 14 | 30 | 22 |
| | **HuMMuS** | GO_Biological_Process_2021 | 1 | 1 | 1 | 1 | 1 | 1 | 1 | 1 | 2 | 4 | 8 | 10 | 14 | 18 | 18 | 25 | 21 | 26 | 29 | 36 | 39 |
| | | GO_Cellular_Component_2021 | 1 | 1 | 1 | 1 | 1 | 1 | 1 | 1 | 2 | 3 | 4 | 7 | 8 | 7 | 8 | 14 | 13 | 19 | 18 | 17 | 30 |
| | | GO_Molecular_Function_2021 | 1 | 1 | 1 | 1 | 1 | 1 | 1 | 1 | 2 | 3 | 3 | 9 | 10 | 10 | 11 | 14 | 21 | 31 | 25 | 36 | 39 |
| | | KEGG_2019_Mouse | 1 | 1 | 1 | 1 | 1 | 1 | 1 | 1 | 2 | 3 | 7 | 8 | 11 | 7 | 6 | 12 | 21 | 22 | 19 | 20 | 25 |
| | | Reactome_2016 | 1 | 1 | 1 | 1 | 1 | 1 | 1 | 1 | 2 | 4 | 4 | 12 | 10 | 15 | 16 | 16 | 20 | 23 | 28 | 26 | 44 |

**Supplementary Table 5. Marker TFs/regulons identified by HuMMuS on the cortical mouse 3-omics dataset.** This table contains the regions and the Wilcoxon test's statistics associated to the identified marker TFs, and the literature supporting their levels of evidence. Different level of literature-based evidence are indicated by colors : green (marker of the specific subpopulation supported by literature), blue (marker of the cortex or similar area supported by literature), yellow (other brain regions / neuron markers supported by literature), orange (expressed in the brain/cortex according to gene expression databases), gray (no evidence)

| gene name | area | Wilcoxon test score / p-values adjusted | associated publication / source * |
|---|---|---|---|
| Tbx1 Tbx10 | Layer 2/3 | 104.65 | loss of Tbx1 disrupts corticogenesis in mice by promoting premature neuronal differentiation |
| | | 0 | https://pubmed.ncbi.nlm.nih.gov/27005988/ |
| Rfx3 | Layer 2/3 | 102.37 | *We [...] identified [...] TFs with more restricted patterns in specific subclasses, such as Rfx3 [...] (in L2/3 IT)* |
| | | 0 | A multimodal cell census and atlas of the mammalian primary motor cortex |
| Vdr (vitamin D) | Layer 2/3 | 102.2 | Expressed in cortical neurons and involved in neurodegeneration |
| | | 0 | https://pubmed.ncbi.nlm.nih.gov/21408608/ |
| Neurog1 | Layer 2/3 | 98.26 | layer II/III neurons of the piriform cortex. |
| | | 0 | https://pubmed.ncbi.nlm.nih.gov/24403153/ |
| Zfp711 | Layer 2/3 | 96.2 | not studied a lot, involved in brain development |
| | | 0 | https://pubmed.ncbi.nlm.nih.gov/20346720/ |
| Pou4f1 Pou4f2 | Layer5a | 81.53 | Expressed in [...] the dorsal column of the mesencephalic and pontine central gray, and the lateral interpeduncular nucleus of the brain |
| | | 0 | https://pubmed.ncbi.nlm.nih.gov/7904822/ |
| Esx1 | Layer5a | 76.13 | expressed highly in midbrain mantle layer (FDR: 4E-4) |
| | | 0 | https://bgee.org/gene/ENSMUSG00000023443?expression=&data_type=IN_SITU |
| Sebox | Layer5a | 71.97 | expressed in cerebral cortex |
| | | 0 | https://www.ncbi.nlm.nih.gov/pmc/articles/PMC16794/ |
| Setbp1 | Layer5a | 70.6 | expressed in ventricular zone |
| | | 0 | https://molecularautism.biomedcentral.com/articles/10.1186/s13229-023-00540-x |
| Pou4f3 | Layer5a | 70.2 | Expressed in brain and DRG |
| | | 0 | https://pubmed.ncbi.nlm.nih.gov/22326227/ |
| Pgr | Layer5 | 79.42 | expressed in substantia niagra |
| | | 0 | https://bgee.org/gene/ENSMUSG00000031870 |
| Nr3C1 | Layer5, Layer6 | 76.48, 75.96 | expressed in median eminence of neurohypophysis |
| | | 0, 0 | https://bgee.org/gene/ENSMUSG00000024431 |
| Mbd2 | Layer5, MGE, Layer5b | 67.94, 60.58, 54.90 | highly expressed in brain |
| | | 0, 0, 0 | https://pubmed.ncbi.nlm.nih.gov/9774669/ |
| Wt1 | Layer5, CGE, MGE | 63.45, 71.84, 60.58 | neurons of DRG and sertolis cells |
| | | 0, 0, 0 | https://pubmed.ncbi.nlm.nih.gov/16467207/ |
| Pbx2 /Pbx4 | Layer5 | 54.88 | regulates patterning f the cerebral cortex in progenitors and post mitotic neurons |
| | | 0 | https://pubmed.ncbi.nlm.nih.gov/26671461/ |
| Olig3 | Claustrum | 29.18 | **Olig3 coordinates the specification of dorsal neurons in the spinal cord** |
| | | 3.54E-187 | https://www.ncbi.nlm.nih.gov/pmc/articles/PMC1065726// |
| Foxn2 | Claustrum | 27.27 | |
| | | 9.80E-164 | |
| Naif1 | Claustrum | 27.09 | |
| | | 1.23E-161 | |
| Dmrtc2 | Claustrum | 26.7 | *DMRT2, DMRTA1/DMRT4, DMRT3 and DMRTA2/DMRT5 are expressed mainly in cortical regions* |
| | | 4.58E-157 | https://www.frontiersin.org/articles/10.3389/fnana.2022.937596/full |
| Cdc5l | Claustrum | 26.61 | |
| | | 5.31E-156 | |
| **Maz** | **CGE** | 73.36 | (1) Expressed in Purkinje cells in the brain (at protein level). / (2) driving neurogenesis |
| | | 0 | (1) https://pubmed.ncbi.nlm.nih.gov/26089202/ (2) https://pubmed.ncbi.nlm.nih.gov/22944911/ |
| Sp2 | CGE | 72.6 | CHECK AGAIN |
| | | 0 | |
| Smad1/Smad5 | CGE, MGE | 71.92, 60.57 | ventricular zone, FDR: 10E-10 |
| | | 0, 0 | https://www.uniprot.org/uniprotkb/P97454/entry#expression |
| Zbtb1 | CGE | 70.19 | |
| | | 0 | |
| Zfp219 | MGE | 60.28 | |
| | | 0 | |
| Klf15 | MGE | 60.05 | |
| | | 0 | |
| Mlxipl | Layer5b | 48.23 | Expressed in the ventricular and intermediate zones of the developing spinal cord of 12.5 dpc embryos. In later embryos expressed in a variety of tissues. |
| | | 0 | https://www.uniprot.org/uniprotkb/Q99MZ3/entry#expression |
| Hoxa1 | Layer5b | 48.19 | Motor neuron axon guidance in development |
| | | 0 | https://pubmed.ncbi.nlm.nih.gov/9367425/ |
| Arntl | Layer5b | 46.21 | constitutively expressed in hypothalamus nucleus suparchiasmatic |
| | | 0 | https://pubmed.ncbi.nlm.nih.gov/11207387/ |
| Mitf | Layer5b | 46 | Microphthalmia-associated transcription factor ensures the elongation of axons and dendrites in the mouse frontal cortex. |
| | | 0 | https://pubmed.ncbi.nlm.nih.gov/27859996/ |
| Mnt | Layer6 | 80.83 | Motor neuron expression |
| | | 0 | https://bgee.org/gene/ENSMUSG00000000282 |
| Zfp282 | Layer6 | 78.68 | |
| | | 0 | |
| Lef1 | Layer6 | 75.96 | deep layers of the cortex (even if more 5 I think?), important for normal development |
| | | 0 | https://www.ncbi.nlm.nih.gov/pmc/articles/PMC3825142/ |
| Tcf7l2 | Layer6 | 74.89 | deep layers of the cortex (even if more 5 I think?), important for normal development |
| | | 0 | https://www.ncbi.nlm.nih.gov/pmc/articles/PMC3825142/ |
